# Jumper Enables Discontinuous Transcript Assembly in Coronaviruses

**DOI:** 10.1101/2021.02.12.431026

**Authors:** Palash Sashittal, Chuanyi Zhang, Jian Peng, Mohammed El-Kebir

## Abstract

Genes in SARS-CoV-2 and, more generally, in viruses in the order of *Nidovirales* are expressed by a process of discontinuous transcription mediated by the viral RNA-dependent RNA polymerase. This process is distinct from alternative splicing in eukaryotes, rendering current transcript assembly methods unsuitable to *Nidovirales* sequencing samples. Here, we introduce the Discontinuous Transcript Assembly problem of finding transcripts 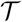 and their abundances **c** given an alignment 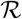 under a maximum likelihood model that accounts for varying transcript lengths. Underpinning our approach is the concept of a segment graph, a directed acyclic graph that, distinct from the splice graph used to characterize alternative splicing, has a unique Hamiltonian path. We provide a compact characterization of solutions as subsets of non-overlapping edges in this graph, enabling the formulation of an efficient mixed integer linear program. We show using simulations that our method, Jumper, drastically outperforms existing methods for classical transcript assembly. On short-read data of SARS-CoV-1 and SARS-CoV-2 samples, we find that Jumper not only identifies canonical transcripts that are part of the reference transcriptome, but also predicts expression of non-canonical transcripts that are well supported by direct evidence from long-read data, presence in multiple, independent samples or a conserved core sequence. Jumper enables detailed analyses of *Nidovirales* transcriptomes.

**Code availability:** Software is available at https://github.com/elkebir-group/Jumper

## 1 Introduction

SARS-CoV-2 is part of the taxonomic order of *Nidovirales*, which comprise enveloped viruses containing a positive-sense, single-stranded RNA genome that encodes for non-structural proteins near the 5’ end as well as structural and accessory proteins near the 3’ end [1]. Since the ribosome starts at the 5’ end, translation of the viral genome only generates the non-structural proteins. Expression of the other genes is achieved by *discontinuous transcription* performed by the viral RNA-dependent RNA polymerase (RdRp) [2], which is present in the non-structural part of the viral genome. Specifically, RdRp may ‘jump’ over contiguous genomic regions, or *segments*, in the viral RNA template. This process results in *discontinuous transcripts* that have matching 5’ and 3’ ends and each correspond to a subsequence of segments ordered as in the reference genome. In a recent paper, Kim *et al*. [3] analyzed sequencing samples showing that SARS-CoV-2 has both canonical discontinuous transcription events that produce an intact 3’ open reading frame (ORF) as well as non-canonical discontinuous transcription events whose role is unclear. To understand the life cycle of *Nidovirales*, it is critical to assemble the complete set 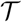 of expressed transcripts and their abundances **c** (Fig. 1A).

**Figure 1:**
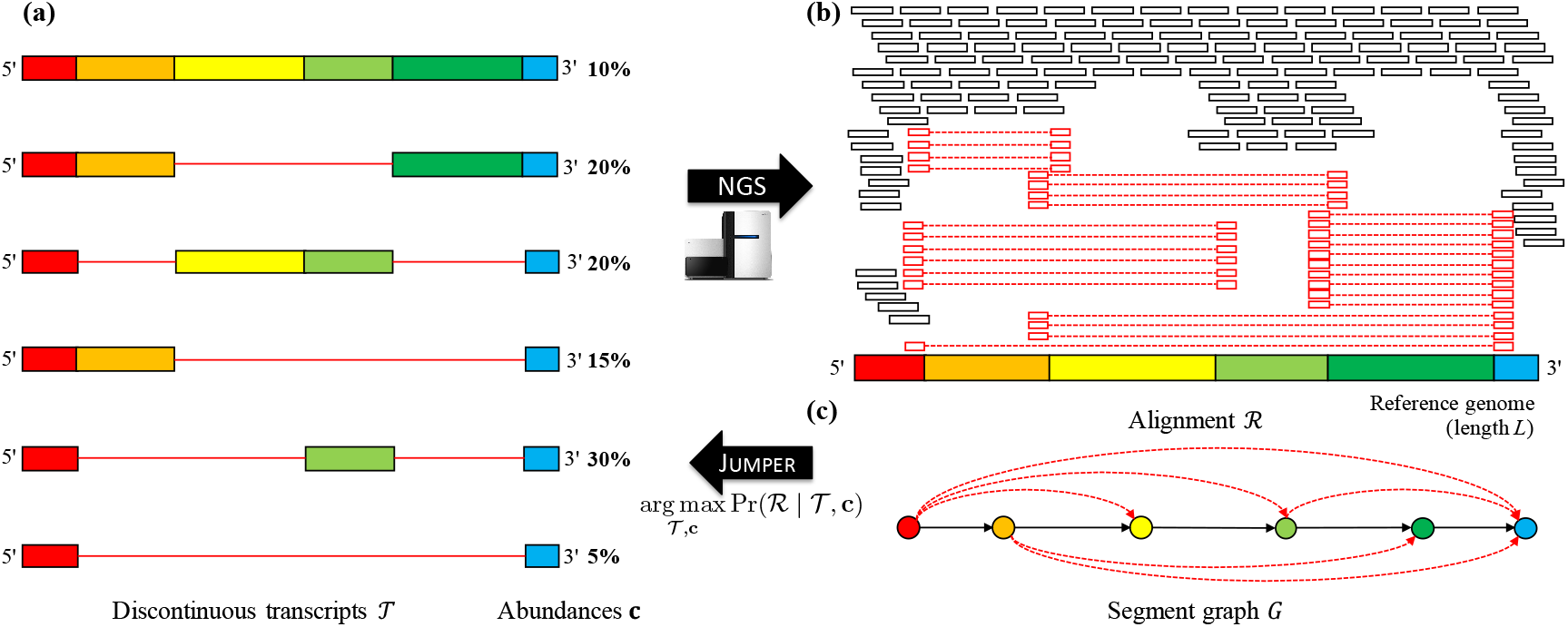
(a) Viruses in the order *Nidovirales* generate a set 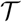 of discontinuous transcripts with varying abundances **c** during infection. (b) Next generation sequencing will produce an alignment 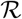 with two types of aligned reads: unphased reads that map to a contiguous genomic region (black) and phased reads that map to distinct genomic regions (red). (c) From 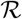 we obtain the segment graph *G*, a directed acyclic graph with a unique Hamiltonian path. Jumper solves the Discontinuous Transcript Assembly to infer 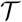 and **c** with maximum likelihood.

There has been a lot of work in methods for transcript assembly from RNA-sequencing data. Briefly, there are two classes of methods, (i) *de novo* assembly methods and (ii) reference-based methods. The main distinction is that reference-based methods require the reference genome as input while *de novo* methods have no such requirement. As such, *de novo* assembly methods [4–8] are useful when the reference genome is unavailable or when the diversity of different species in the sample is too large. On the other hand, reference-based methods [9–13] are able to achieve higher accuracy as the reference genome provides a scaffold on which to align sequencing reads. Importantly, current methods from both classes are designed to assemble transcripts in eukaryotes, where transcripts expressed by a gene may differ in their composition due to alternative splicing. Alternative splicing is predominantly mediated by the spliceosome, which results in the removal of introns, defined by conserved splice sites, from the transcribed pre-mRNA. This is a distinct process compared to discontinuous transcription in *Nidovirales*, which is mediated by viral RdRp, which results in the removal of contiguous segments due to jumps of the RdRp at locations that are not fully characterized. As such, current methods for transcript assembly that have been mainly designed for use in eukaryotes are not optimized for transcript assembly in *Nidovirales*.

In this study, we introduce the Discontinuous Transcript Assembly problem of finding discontinuous transcripts 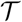 and their abundances **c** (Fig. 1b) given an alignment 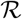. Underpinning our approach is the concept of a segment graph (Fig. 1c), a directed acyclic graph that, distinct from the splice graph used to characterize alternative splicing, has a unique Hamiltonian path. We characterize solutions as subsets of non-overlapping edges in this graph, enabling the formulation of an efficient mixed integer linear program while accounting for paired-end read information. Using simulations, we show that our method, Jumper, drastically outperforms Scallop [9] and String Tie [10], existing methods for transcript assembly. In real data [3], we run Jumper on paired-end short-read data of virus infected Vero cells and use long-read data of the same sample for validation. We find that Jumper not only identifies canonical transcripts that are part of the reference transcriptome, but also predict expression of non-canonical transcripts that are well supported by long-read data. Similarly, Jumper identifies canonical and non-canonical transcripts in SARS-CoV-1 samples [14]. In summary, Jumper enables detailed *de novo* analyses of *Nidovirales* transcriptomes.

## 2 Preliminaries and Problem Statement

We define discontinuous transcripts as follows.

### Definition 1.

Given a reference genome, a *discontinuous transcript T* is a sequence **v**_1_, … , **v**_|*T*|_ of segments where (i) each segment corresponds to a contiguous region in the reference genome, (ii) segment **v**_*i*_ precedes segment **v**_*i*+1_ in the reference genome for all *i* ∈ {1, … , |*T*| − 1}, (iii) segment **v**_1_ contains the 5’ end of the reference genome and (iv) segment **v**_|*T*|_ contains the 3’ end of the reference genome.

In the literature, discontinuous transcripts that differ from the genomic transcript *T*_0_ are called *subgenomic transcripts*, which correspond to subgenomic RNAs (sgRNAs) [3]. Transcripts 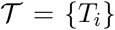 occur in *abundances* **c** = [*c_i_*] where *c_i_* ≥ 0 is the relative abundance of transcript *T_i_* such that 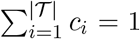. While next-generation sequencing technologies provide high coverage of the viral genome of length *L* of about 10 to 30 Kbp, they are limited to short reads with fixed length *ℓ* ranging from 100 to 400 bp. For ease of exposition, we describe the formulation in context of single-end reads, but in practice we use the paired-end information if it is available. As *ℓ* ≪ *L*, the identity of the transcript of origin for a given read is ambiguous. Therefore we need to use computational methods to reconstruct the transcripts and their abundances from the sequencing reads. Specifically, given a *Nidovirales* reference genome of length *L* and reads of a fixed length *ℓ*, we use a splice-aware aligner such as `STAR` [15] to obtain an alignment 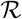. This alignment provides information about the abundance **c** and composition of the underlying transcripts 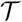 in the following two ways. First, the *depth*, or the number of reads along the genome is informative for quantifying the abundance **c** of the transcripts. Second, the composition 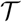 of the transcripts is embedded in *phasing reads*, which are reads that align to multiple distinct regions in the reference genome (Fig. 1B).

To make the relationship between 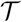, **c** and 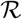 clear, we introduce the *segment graph G*, which is obtained from the phasing reads in a alignment 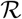. As mentioned, each phasing read 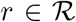 maps to *q* ≥ 2 distinct regions in the reference genome. Each pair of regions that are adjacent in the phasing read are separated by two positions *v, w* (where *w* − *v* ≥ 2) in the reference genome called *junctions*. Thus, each phasing read contributes 2*q* − 2 junctions. The collective set of junctions contributed by all phasing reads in 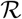 in combination with positions {1*, L*} induces a partition of the reference genome into closed intervals [*v*^−^*, v*^+^] of junctions that are consecutive in the reference genome (*i.e.* there exists no other junction that occurs in between *v*^−^ and *v*^+^). The resulting set of segments equals the node set *V* of segment graph *G* (Fig 2a). The edge set *E* of segment graph *G* is composed of continuous edges *E*^→^ and discontinuous edges *E^↷^*. Continuous edges *E*^→^ are composed of ordered pairs (**v** = [*v*^−^*, v*^+^], **w** = [*w*^−^*, w*^+^]) of nodes that correspond to segments that are adjacent in the reference genome, *i.e.* where *v*^+^ = *w*^−^. On the other hand, discontinuous edges *E^↷^* are composed of ordered pairs (**v** = [*v*^−^*, v*^+^], **w** = [*w*^−^*, w*^+^]) of nodes that corresponds to segments that are adjacent in at least one phasing read in 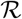 but not adjacent in the reference genome (*i.e. w*^−^ − *v*^+^ ≥ 2). Fig. 1c shows an example of a segment graph.

**Figure 2:**
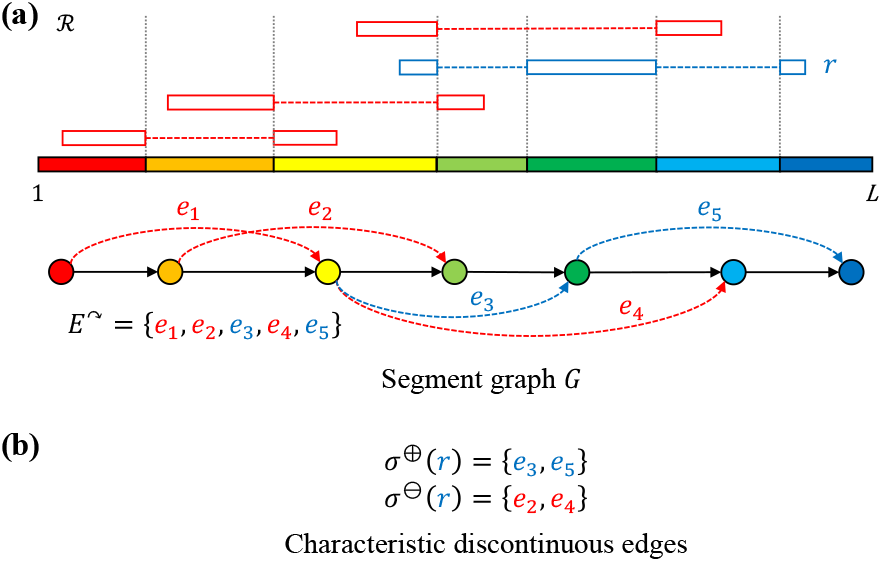
(a) Phasing reads in an alignment 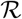 define a set of junctions, which in turn define the segment graph *G*. (b) Each phasing read has characteristic discontinuous edges indicating the set *σ*^⊕^ of discontinuous edges present in the read as well as conflicting/overlapping discontinuous edges *σ*^⊖^. Here, phasing read Characteristic discontinuous edges *r* (blue), has *σ*^⊕^(*r*) = {*e*_3_ , *e*_5_} and *σ*^⊖^(*r*) = {*e*_2_*, e*_4_}. Note that *e*_1_ is not included in *σ*^⊖^(*r*) as it does not overlap with *π*(*r*) = {*e*_3_*, e*_5_}.

### Definition 2.

Given an alignment 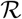, the corresponding *segment graph G* = (*V, E*^→^ ∪ *E^↷^*) is a directed graph whose node set *V* equals the set of segments induced by the junctions of phasing reads in 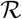 and whose edge set *E* = *E*^→^ ∪ *E^↷^* is composed of edges (**v** = [*v*^−^*, v*^+^], **w** = [*w*^−^*, w*^+^]) that are either continuous, *i.e. v*^+^ = *w*^−^, or discontinuous, *i.e. w*^−^ − *v*^+^ ≥ 2 and there exists a phasing read where junctions *v*^+^ and *w*^−^ are adjacent.

We note that the segment graph *G* is closely related to the splice graph used in regular transcript assembly where transcripts correspond to varying sequences of exons due to alternative splicing. The key difference, however, is that an alignment 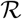 generated from reads obtained from discontinuous transcripts induces a segment graph *G* that is a directed acylic graph (DAG) with a unique Hamiltonian path. This is because, as stated in Definition 1, discontinuous transcripts 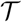 have matching 5’ and 3’ ends, and, although their comprising segments may vary, their order follows the reference genome.

### Observation 1.

Segment graph *G* is a directed acyclic graph with a unique Hamiltonian path.

The unique Hamiltonian path of *G* corresponds to the sequence of continuous edges *E*^→^. This path corresponds to the whole viral genome which is generated by the RdRp during the replication step [2]. Moreover, by the above observation, *G* has a unique source node **s** and sink node **t**. Importantly, each transcript 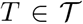 that is compatible with an alignment 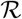 corresponds to an **s** − **t** path *π*(*T*) in *G*. Here, a *path π* is a subset of edges *E* that can be ordered (**v**_1_, **w**_1_), … , (**v**_|*π*|_, **w**_|*π*|_) such that **w**_*i*_ = **v**_*i*+1_ for all *i* ∈ [|*π*| − 1] = {1, … , |*π*| − 1}. While splice graphs are DAGs and typically have a unique source and sink node as well, they do not necessarily contain a Hamiltonian path [9, 16–18].

Our goal is to find a set 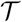 of transcripts and their abundances **c** that maximize the posterior probability

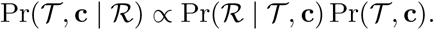

Under an uninformative, flat prior, this is equivalent to maximizing the probability 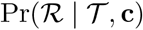. We use the segment graph *G* to compute the probability 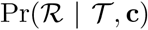 of observing an alignment 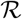 given transcripts 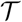 and abundances **c**. We follow the generative model which has been extensively used for transcription quantification [19–21]. The notations used in this paper best resemble the formulation described in [18]. Let 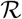 be composed of reads be {*r*_1_, … , *r_n_*} and the set 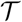 of transcripts be 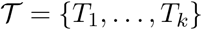 with lengths *L*_1_, … , *L_k_* and abundances **c** = [*c*_1_, … , *c_k_*]. In line with current literature, reads 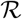 are generated independently from transcripts 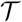 with abundances **c**. Further, we must marginalize over the set of transcripts 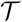 as the transcript of origin of any given read is typically unknown, since *ℓ* ≪ *L*. Moreover, we assume that the fixed read length *ℓ* is much smaller than the length *L_i_* of any transcript *T_i_*. As such, we that 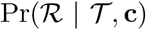 equals

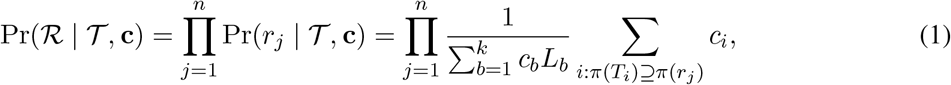

where *π*(*T*) ⊆ *E* is the **s** − **t** path corresponding to transcript *T* and *π*(*r*) ⊆ *E* is the path induced by the ordered sequence of segments (or nodes of *G*) spanned by read *r*. By construction, *π*(*T*) ⊇ *π*(*r*) is a necessary condition for transcript *T* to be a candidate transcript of origin of read *r*. Appendix A gives the derivation of the above equation (Eq. (1)). Our goal is to find 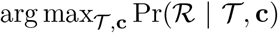, leading to the following problem.

### Problem 1

(Discontinuous Transcript Assembly (DTA)). Given alignment 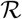 and integer *k*, find discontinuous transcripts 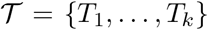 and abundances **c** = [*c*_1_, … , *c_k_*] such that (i) each transcript 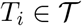 is an **s** − **t** path in segment graph *G*, and (ii) 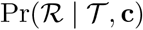 is maximum.

## 3 Combinatorial Characterization of Solutions

Eq. (1) defines the probability 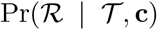 in terms of the observed reads *r* and their induced paths *π*(*r*) ⊆ *E*(*G*) in the segment graph *G*. The authors in [18] use this characterization of reads as paths in a general *splice* graph to account for ambiguity in the transcript of origin for the reads. For a general splice graph, such a characterization is required to capture all the possible observed reads. However, in our setting, where the segment graph *G* is a DAG with a unique Hamiltonian path, it is possible to describe each read and each transcript *uniquely* in a more concise form. Each path in the segment graph is characterized by a set of *non-overlapping* discontinuous edges. To describe this, we introduce the following definition.

### Definition 3.

Two edges (**v** = [*v*^−^*, v*^+^], **w** = [*w*^−^*, w*^+^]) and (**x**= [*x*^−^*, x*^+^], **y** = [*y*^−^*, y*^+^]) of *G overlap* if the open intervals (*v*^+^*, w*^−^) and (*x*^+^*, y*^−^) intersect, *i.e.* (*v*^+^*, w*^−^) ∩ (*x*^+^*, y*^−^) ≠ ∅.

For any transcript *T* corresponding to an **s** − **t** path in *G*, for which we are only given its discontinuous edges *σ*(*T*), the continuous edges of *T* are uniquely determined by *G* and *σ*(*T*). That is, the continuous edges of *T* equal precisely the subset of continuous edges *E*^→^ that do *not* overlap with any of the discontinuous edges in *σ*(*T*). Conversely, given an **s** − **t** path *π*(*T*) of *G* the corresponding set of discontinuous edges is given by *σ*(*T*) = *π*(*T*) ∩ *E^↷^*. Thus, we have the following proposition with the proof in Appendix B.1.

### Proposition 1.

There is a bijection between subsets of discontinuous edges that are pairwise non-overlapping and **s** − **t** paths in *G*.

In a similar vein, rather than characterizing a read *r* by its induced path *π*(*r*) ⊆ *E* in the segment graph, we characterize a read *r* by a pair (*σ*^⊕^(*r*), *σ*^⊖^(*r*)) of *characteristic discontinuous edges*. Here, *σ*^⊕^(*r*) is the set of discontinuous edges that must be present in any transcript that could generate read *r*, *i.e. σ*^⊕^(*r*) = *π*(*r*) ∩ *E^↷^*. Conversely, *σ*^⊖^(*r*) is the set of discontinuous edges that must be absent in any transcript that could generate read *r* due to the unidirectional nature of RdRp transcription. Thus, the set *σ*^⊖^(*r*) consists of discontinuous edges *E^↷^* \ *σ*^⊕^ that overlap with any edge in *π*(*r*). Clearly, while *σ*^⊕^(*r*) ∩ *σ*^⊖^(*r*) = ∅, it need not hold that *σ*^⊕^(*r*) ∪ *σ*^⊖^(*r*) equals *E^↷^* (see Fig. 2b). Formally, we define (*σ*^⊕^(*r*), *σ*^⊖^(*r*)) as follows.

### Definition 4.

The *characteristic discontinuous edges* of a read *r* are a pair (*σ*^⊕^(*r*), *σ*^⊖^(*r*)) where *σ*^⊕^(*r*) is the set of discontinuous edges present in read *r*, *i.e. σ*^⊕^(*r*) = *π*(*r*) ∩ *σ*^⊖^, and 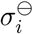 is the set of discontinuous edges (**v** = [*v*^−^*, v*^+^], **w** = [*w*^−^*, w*^+^]) ∈ *E^↷^* \ *σ*^⊕^(*r*) that *overlaps* with an edge (**x** = [*x*^−^*, x*^+^], **y** = [*y*^−^*, y*^+^]) in *π*(*r*).

We have the following result with the proof given in Appendix B.1.

### Proposition 2.

Let *G* be a segment graph, *T* be a transcript and *r* be a read. Then, *π*(*T*) ⊇ *π*(*r*) if and only if *σ*(*T*) ⊇ *σ*^⊕^(*r*) and *σ*(*T*) ∩ *σ*^⊖^(*r*) = ∅.

Hence, we may rewrite the likelihood 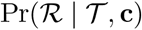 as

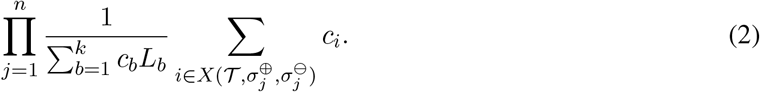

where 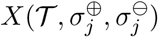 be the subset of indices *i* corresponding to transcripts 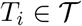 where 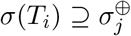 and 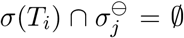. Note that the only difference between Eq. (2) and the formulation in Eq. (1) is the way that the candidate transcripts of origin for a given read are described. In Eq. (1), they are described as paths in the splice graph wheres in Eq. (2), they are described by sets of pairwise non-overlapping discontinuous edges in the segment graph. This leads to the following theorem.

### Theorem 1.

For any alignment 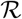, transcripts 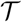 and abundances **c**, Equations (1) and (2) are identical.

Although we have described the formulation for single-end reads, this characterization is applicable to paired-end and even synthetic long reads. Moreover, our implementation provides support for both single-end and paired-end read samples with a fixed read length. The above characterization using discontinuous edges allows us to reduce the number of terms in the likelihood function since multiple reads can be characterized by the same characteristic discontinuous edges. We describe this in detail in Section 4.

## 4 Methods

To solve the DTA problem, we use the results of Section 3 to write a more concise form of the likelihood. Specifically, let 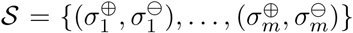 be the set of characteristic discontinuous edges generated by the reads in alignment 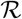. Let **d** = {*d*_1_, · · · , *d_m_*} be the number of reads that map to each pair in 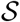. Using that reads *r* with identical characteristic discontinuous edges (*σ*^⊕^(*r*), *σ*^⊖^(*r*)) have identical probabilities 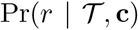, we obtain the following mathematical program for the log-likelihood log 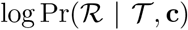 (see Appendix A for derivation).

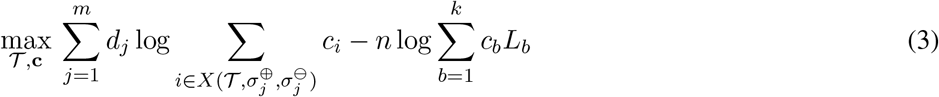

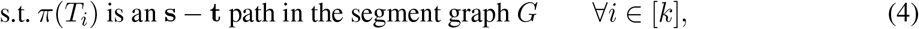

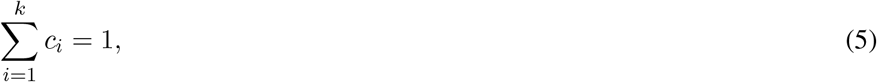

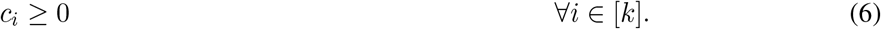

Observe that the first sum (over reads) is concave and the second sum (over transcripts) is convex. Since we are maximizing, our objective function would ideally be concave. In Appendix B.1, we prove the following lemma, which enables us to remove the second term using a scaling factor for the relative abundances **c** that does not alter the solution space.

### Lemma 1.

Let *D* > 0 be a constant, 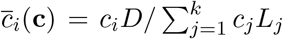 and 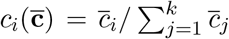 for all *i* ∈ [*k*]. Then, 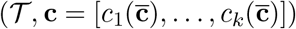 is an optimal solution for (3)-(6) if and only if 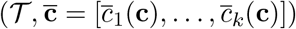 is an optimal solution for

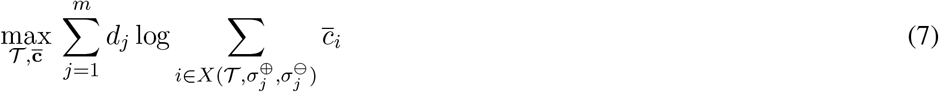

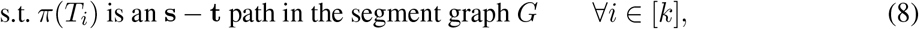

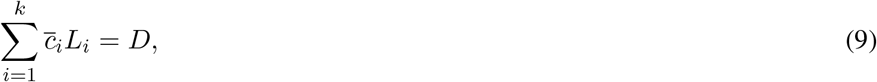

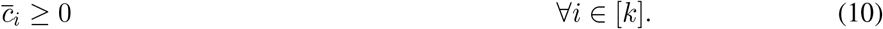

We formulate the mathematical program given in Lemma 1 as a mixed integer linear program. More specifically, we encode (i) the composition of each transcript *T_i_* as a set *σ*(*T_i_*) of non-overlapping discontinuous edges, (ii) the abundance *c_i_* and length *L_i_* of each transcript *T_i_*, (iii) the total abundance 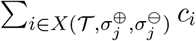 of transcripts supported by characteristic discontinuous edges 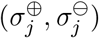, and (iv) a piece-wise linear approximation of the log function using a user-specified number *h* of breakpoints. We will describe (i) and (ii) in the following and refer to Appendix B.2 for (iii) and (iv).

### Transcript composition

We begin modeling (8), which states that each transcript *T_i_* must correspond to an **s** − **t** path in the segment graph *G*. Using Proposition 1, we introduce binary variables 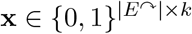 to encode the presence of discontinuous edges in each of the *k* **s** − **t** paths corresponding to the *k* transcripts in 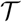. For any discontinuous edge *e* = (**v** = [*v*^−^*, v*^+^], **w** = [*w*^−^*, w*^+^]), let *I*(*e*) denote the open interval (*v*^+^*, w*^−^) between the two segments **v** and **w**. By Proposition 1, it must hold that *I*(*e*) ∩ *I*(*e*′) = ∅ for any two distinct discontinuous edges *e* and *e*′ assigned to the same transcript. To encode this, we impose

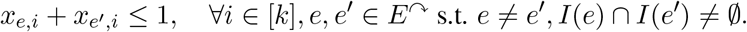

### Transcript abundance and length

We introduce non-negative continuous variables **c** = [*c*_1_*, … , c_k_*] that encode the abundance of the *k* transcripts. The scale of these abundances depends on the choice of *D*. We choose *D* = *ℓ*\* where *ℓ*\* is the length of the shortest **s** − **t** path in the segment graph *G*. Substituting *D* = *ℓ*\* into (9) yields 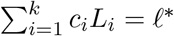.

Since 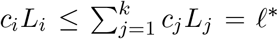 and *L_i_* ≥ *ℓ*\*, we have that *c_i_* ≤ 1. To model the product *c_i_L_i_* of the length *L_i_* of a transcript *T_i_* and its abundance *c_i_*, we focus on individual discontinuous edges *e*. For any discontinuous edge *e* = (**v** = [*v*^−^*, v*^+^], **w** = [*w*^−^*, w*^+^]), let *L*(*e*) = *w*^−^ − *v*^+^ be the length of the interval. Observe that

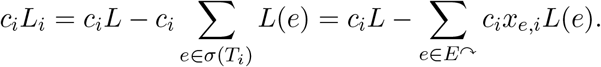

We introduce continuous variables *z_e_* ∈ [0, 1]^*k*^ and encode the product *z_e,i_* = *c_i_x_e,i_* for all *e* ∈ *E^↷^* as

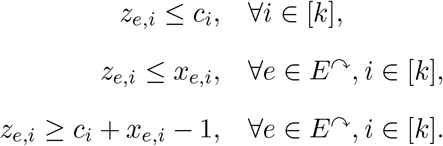

Therefore, we may represent 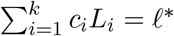 as

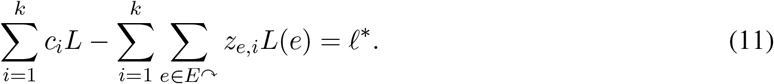

The resulting formulation has *O*(|*E^↷^*|*k* +|*E^↷^*|*m*+*mh*) variables, where *h* is the user-specified number of breakpoints used in the piecewise linear approximation of the log function. This number includes |*E^↷^*|*k* binary variables. The number of constraints is *O*(*k*|*E*|^2^ + |*E*|*km*).

### Progressive heuristic

In practice, the number of discontinuous edges in the segment graph is inflated due to ambiguity in the exact location at which the RdRp jumps as well as sequencing and alignment errors. This leads to large number of binary variables in our MILP (we have *k* · |*E^↷^*| binary variables) which can make the MILP intractable. In order to approximately solve the problem with large values of *k*, we implement a progressive heuristic. Our heuristic takes as input the alignment 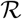 and an integer *k*, which is the maximum number of transcripts in the solution. At each iteration *p* ≤ *k*, we are given a set 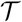 of *p* − 1 previously computed transcripts and seek a new transcript *T*′ by solving the MILP (Eq. (27)-(31)) using function solveILP with additional constraints to fix the values of the variables that encode the presence/absence of discontinuous edges for the transcripts in 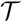. The resulting reduction in number of binary variables from |*E^↷^*|*k* to |*E^↷^*| improves the running time of the MILP. As an additional optimization, we re-estimate the abundances of a new set 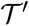 of transcripts. This set contains all transcripts in 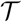 as well additional transcripts corresponding to all possible subsets of discontinuous edges *σ*(*T*′) of the newly identified transcript *T*′, identified by the function Expand. We solve a linear programming (Eq. (32)–(34)) with function SolveLP to re-estimate the abundances **c**′ of 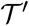, retaining only the top *p* transcripts *T_i_* from 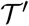 with the largest abundances *c_i_L_i_*. We terminate upon convergence, *i.e.* if 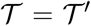, or if the number *p* of iterations reaches the number *k*. Algorithm 1 provides the pseudo code of the progressive heuristic implemented in Jumper. The details of the subproblems SolveILP and SolveLP are given in Appendix B.3.

### Implementation details

Matching core sequences that mediate the discontinuous transcription by RdRp lead to ambiguity in precise location of breakpoint during alignment of spliced reads. Therefore, in practice we observe multiple discontinuous edges with closely spaced 5’ and 3’ breakpoints. Moreover, false positive discontinuous edges are introduced due to sequencing and alingment errors. We use a threshold on the number of spliced reads supporting a discontinuous edge to filter false positive edges with low support. This parameter can also be used to reduce computational burden and focus on the highly expressed transcripts in the sample. A discussion on the choice of the thresholding parameter is provided in Section B.4. Jumper is implemented in Python 3 using `Gurobi` [22] (version 9.0.3) to solve the MILP and `pysam` [23] for reading and processing the input BAM file. Jumper is available at https://github.com/elkebir-group/Jumper.

**Algorithm 1:**
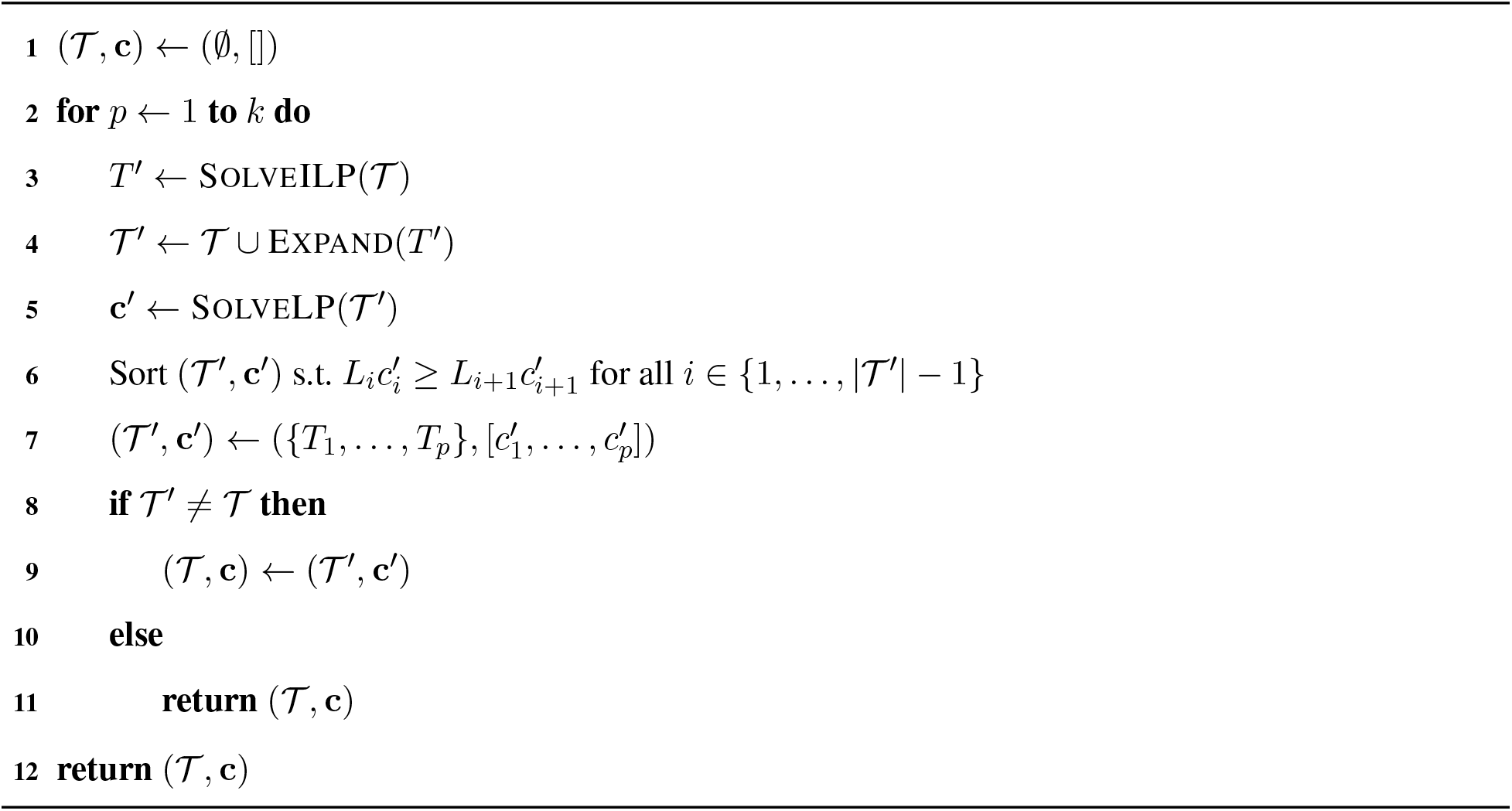
Jumper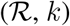

## 5 Results

Before we describe our results for coronavirus samples, we establish some terminology that will be used in this section. A discontinuous edge (**v** = [*v*^−^*, v*^+^], **w** = [*w*^−^*, w*^+^]) is *canonical* provided its 5’ junction *v*^+^ occurs in the transcription regulating leader sequence (TRS-L), *i.e.* between positions 55 and 85 and the first occurrence of ‘AUG’ downstream of the 3’ junction *w*^−^ position coincides with the start codon of a known ORF, otherwise the discontinuous edge is called *non-canonical*. In a similar vein, a transcript is *canonical* if it contains at most one canonical and no non-canonical discontinuous edges, otherwise the transcript is *non-canonical*. We run our experiments on a server with two 2.6 GHz CPUs and 512 GB of RAM. We follow the approach of Scallop and use Salmon [20] to accurately re-estimate the abundance of the transcripts assembled by Jumper.

### 5.1 Simulations

We begin our simulations with a segment graph *G* obtained from a short-read sample (SRR11409417). Following Kim *et al.* [3], we used `fastp` to trim short reads (trimming parameter set to 10 nucleotides), which were input to `STAR` run in two-pass mode yielding an alignment 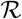. Fig. 3a shows the sashimi plot of the *canonical* and the *non-canonical* discontinuous edges (mappings) supported by the reads in the sample. From 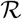, we obtained *G* by only including discontinuous edges supported by at least 20 reads. The segment graph for this sample has |*V* | = 39 nodes and |*E*| = 67 edges, which include |*E^↷^*| = 29 discontinuous edges and |*E*^→^| = 38 continuous edges. The former are subdivided into 14 canonical discontinuous edges that produce a known ORF and 15 non-canonical discontinuous edges. The next step of our simulation pipeline is to generate transcripts 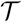 and their abundances **c** for the given segment graph. We used the negative-sense discontinuous transcription model (described in Appendix C.1). Upon generating the transcripts, we simulated the generation and sequencing of RNA-seq data, and aligned the simulated reads using `STAR` [15]. We generated 5 independent pairs 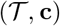 of transcripts and abundances under the negative-sense discontinuous transcription model. Fig. 3b shows the number of transcripts generated from each simulation using the negative-sense discontinuous transcription model. For each pair 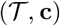 we ran 5 paired-end short read sequencing simulations using `polyester` [24]. Therefore, we generated a total of 5 × 5 = 25 simulated instances. Details are provided in Appendix C.1. Note that our method Jumper does *not* use the negative-sense discontinuous transcription model to infer the viral transcripts from the simulated data.

**Figure 3:**
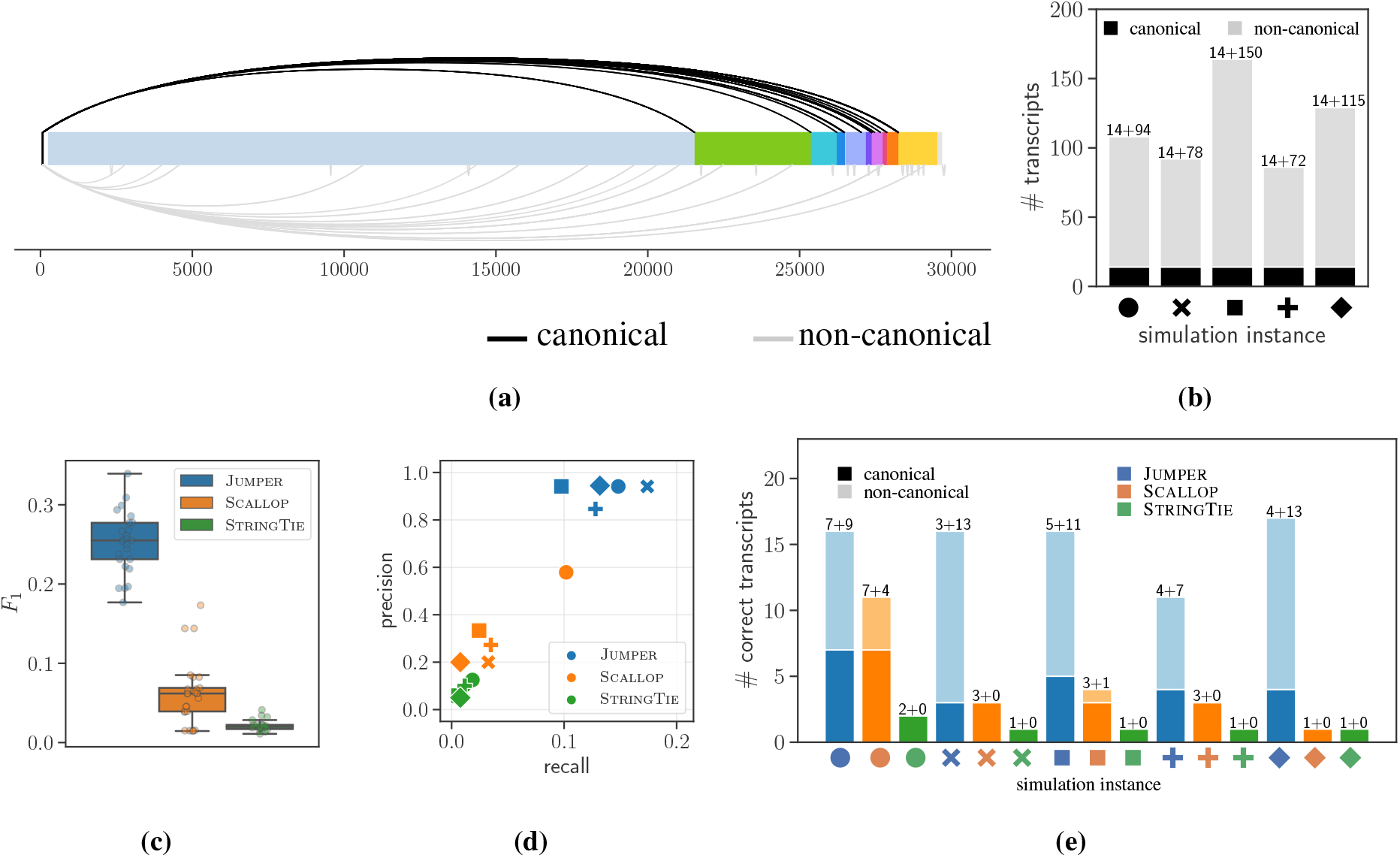
Jumper consistently outperforms Scallop and StringTie in reconstruction of viral transcripts from simulated SARS-CoV-2 sequencing data. (a) Sashimi plot showing the canonical (black) and non-canonical (gray) discontinuous mappings supported by reads in short-read sample SRR11409417. (b) Number of canonical and non-canonical transcripts for 5 simulation instances of 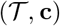 generated under the negative-sense discontinuous transcription model. (c) *F*_1_ score of the three methods (Jumper, Scallop and StringTie) for all the 25 simulated instances under the negative-sense discontinuous transcription model. (d) Precision and recall values of the three methods with one of sequencing experiment for each simulated instance of 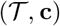 under the negative-sense discontinuous transcription model as input. (e) Total number of canonical and non-canonical transcripts recalled by the three methods for the simulated instances shown in panel (d).

We compare the performance of our method Jumper with two other reference-based transcript assembly methods, Scallop and StringTie. All the methods were run with their default parameters. To avoid including false-positive discontinuous edges, Jumper discards discontinuous edges with support less than 100 reads (Λ = 100) from the segment graph. The reader is referred to Appendix B.4 for a discussion on the choice of the Λ parameter and comparison with other methods. We compare the transcripts predicted by the three methods to the ground truth transcripts. Specifically, a predicted transcript is *correct* (or matches the ground truth) if there exists a transcript in the ground truth whose junction positions match the predicted junctions positions within a tolerance of 10 nucleotides.

Fig. 3c shows the *F*_1_ score (harmonic mean of recall and precision) of the three methods for all the simulated sequencing data instances under the negative-sense discontinuous transcription model. Jumper achieves a much higher *F*_1_ score compared to Scallop and StringTie. For the 25 simulation instances, Jumper has a median *F*_1_ score of 0.255 (range [0.176, 0.339]) while Scallop has a median *F*_1_ score of 0.062 (with range [0.0145, 0.173]) and StringTie has a median *F*_1_ score of 0.01904 (with range [0.0114, 0.0412]). Fig. C2 shows that this trend holds for recall and precision values and our method has a run-time comparable to the existing methods. To investigate the effect of threshold parameter Λ on the performance of Jumper, we ran our method on the simulated instances with Λ ∈ {10, 50, 100, 200}. Fig. C3 shows that Jumper outperforms Scallop and StringTie for all values of Λ, although it incurs significantly more runtime for Λ = 10. Thus, we find that Jumper consistently performs better than Scallop and StringTie.

To better understand the tradeoff between precision and recall, we zoom in one five distinct pairs 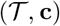 of simulation instance. Fig. 3d shows the precision and recall values achieved by each method for each of these five simulation instances, demonstrating that Jumper consistently outperforms both Scallop and StringTie. On average, Jumper recalls 5 times more transcripts than Scallop and 11 times more transcripts than StringTie while also having higher precision in all simulated cases. Fig. C4 shows that all three methods produce similar precision and recall values for different sequencing replicates of the same simulated instance of 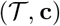, demonstrating consistency in results. Finally, Fig. 3e shows the number of canonical and non-canonical transcripts generated by the three methods for each simulated instance that match the ground truth. In summary, we found that Jumper correctly predicts higher number of both canonical and non-canonical transcripts compared to Scallop and StringTie for all the simulated instances. We observe similar trends on simulated instances of a human gene (see Appendix C.4). The results from all the simulations are summarized in Table C1.

### 5.2 Transcript assembly in SARS-CoV-2

In a recent paper, Kim *et al.* [3] explored the transcriptomic architecture of SARS-CoV-2 by performing short-read as well as long-read sequencing of Vero cells infected by the virus. The authors used oligo(dT) amplification, which targets the poly(A) tail at the 3’ end of messenger RNAs, thus limiting positional bias that would occur when using SARS-CoV-2 specific primers [25, 26]. Subsequently, the authors aligned the resulting reads using splice-aware aligners, *i.e.* `STAR` [15] for the short-read sample (median depth of 1763) and `minimap2` [27] for the long-read sample (median depth of 6707 and mean length of 2875 bp). For both complementary sequencing techniques, the authors observed phasing reads that were indicative of canonical as well as non-canonical transcription events. While the authors quantified the fraction of phasing reads supporting each discontinuous transcription event, they did not attempt to assemble complete transcripts.

We use Jumper to reconstruct the SARS-CoV-2 transcriptome of the short-read sequencing sample using the BAM file obtained by running Kim *et al.*’s pipeline [3], followed by running Salmon to identify precise transcript abundances. We note that running Scallop on the short-read data resulted in only a single, complete canonical transcript (corresponding to ‘N’) but required subsampling of the BAM file (to 20%) due to memory constraints, whereas StringTie produced two incomplete transcripts (‘ORF3a’ and and a non-canonical transcript with low support). We build a segment graph with |*V* | = 59 nodes and |*E*| = 93 edges comprising of |*E^↷^*| = 35 most abundant discontinuous edges, 18 of which canonical and 17 non-canonical (Fig. 4a). Jumper identifies 33 transcripts, 17 of which have an abundance of at least 0.001 as determined by Salmon. We focus our attention to these 17 transcripts (Fig. 4b). A subset of 8 transcripts are canonical, thus containing at most one discontinuous edge with the 5’ junction in TRS-L and the first ATG downstream of the 3’ junction coinciding with the start codon of a known ORF. These canonical transcripts correspond to ORF1ab, ORF3a, E, M, ORF7a, ORF7b, ORF8, N. In particular, ORF1ab (abundance of 0.008) corresponds to the complete viral genome, necessary for viral replication. Notably, ORF10 is the only missing ORF in the identified transcriptome, which is in line with previous studies [3, 28] that did not find evidence for active transcription of ORF10.

**Figure 4:**
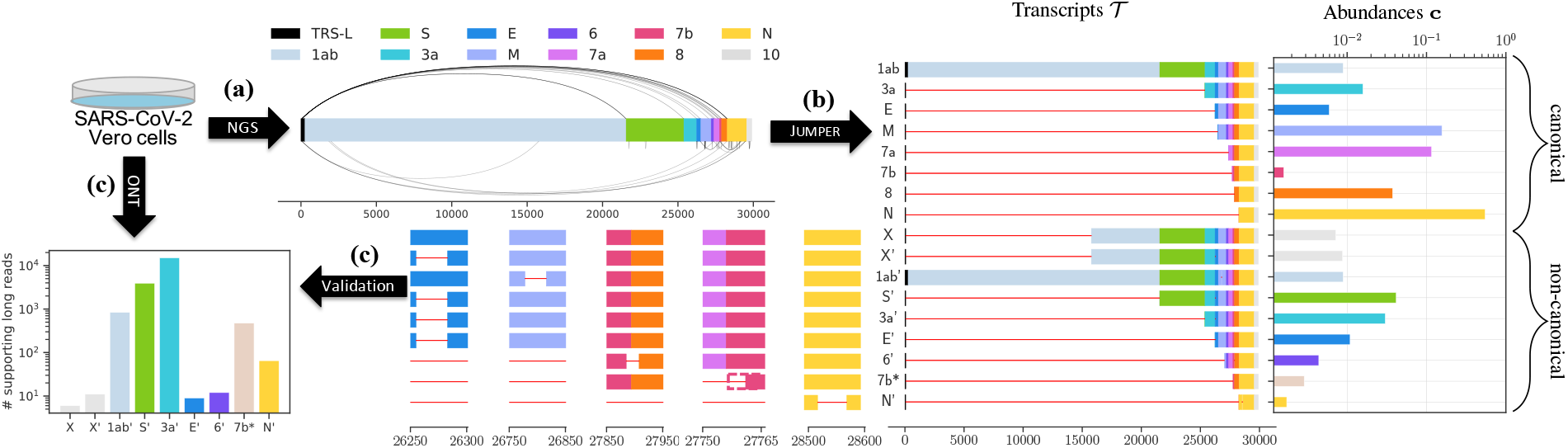
Using short-read data of SARS-CoV-2 infected Vero cells [3], Jumper identifies canonical and non-canonical transcripts that are well supported by long-read sequences of the same sample. (a) The segment graph for the short-read data contains both canonical (above) and non-canonical (below) edges. (b) Jumper assembles 8 canonical transcripts and 9 non-canonical transcripts and estimates their abundances with zoomed-in view of the non-canonical transcripts X, X’, 1ab’, S’, 3a’, E’, 6’, 7b* and N’. (c) All non-canonical transcripts predicted by Jumper are well supported by long-read data. (NGS: Next Generation Sequencing; ONT: Oxford Nanopore Technologies)

As mentioned, Jumper inferred 9 non-canonical transcripts, denoted as X, X’, 1ab’, S’, 3a’, 6’, E’, 7b* and N’. Among these, transcripts 1ab’, S’, 3a’ and 6’ encode for the 1ab polypeptide, spike protein S and accessory protein 3a and accessory protein 6, respectively. Transcripts X and X’ both contain the discontinuous edge going from position 68 to 15774, with the latter containing an additional discontinuous edge from position 26256 to 26284. The 5’ end of the common discontinuous edge occurs within TRS-L, whereas the 3’ end occurs in the middle of ORF1b but is out of frame with respect to the starting position of ORF1b (13468). Specifically, the start codon ‘ATG’ downstream of the 3’ end is located at position 15812 and occurs within nsp12 (RdRp) and the first stop codon is located at position 15896, encoding for a peptide sequence of 28 amino acids. Interestingly, when we examine the reference genome, we observe matching sequences “GAACTTTAA” near the 5’ and 3’ junctions of the discontinuous edge common to X and X’, possibly explaining why the viral RdRp generated this jump (Fig. C5a,b). Strikingly, both matching sequences are conserved within the *Sarbecovirus* subgenus but not in other subgenera of the *Betacoronavirus* genus (Fig. C5a,c). To further corroborate this transcript, we examined short and long-read SARS-CoV-2 sequencing samples from the NCBI Sequence Read Archive (SRA). Specifically, we looked for the presence of reads potentially originating from transcript X focusing on high-quality samples with 100 or more leader-spanning reads (reads whose 5’ end intersects with the TRS-L region). A read *r supports* a transcript *T* if the discontinuous edges of *r* exactly match those of *T* , *i.e. π*(*r*) ⊆ *π*(*T*) and |*σ*^⊕^(*r*)| = |*σ*(*T*)| (Fig. C6). We find ample support for transcript X in both short and long-read samples on SRA, with 100 out of 351 short-read samples and 81 out of 653 long-read samples having more than 0.001 leader-spanning reads supporting transcript X (Fig. C7). We note that although this discontinuous transcription event was also observed in [28], the authors found no evidence of this transcript leading to protein product in the ribo-seq data. Further research into a potentially regulatory function of this transcript is required.

As stated, the difference between transcripts X and X’ is that the latter includes an additional discontinuous edge, corresponding to a short jump of | 27 nucleotides between positions 26256 and 26284. This is an in frame deletion inside ORF E, resulting in the loss of 9 amino acids that span the N-terminal domain (4 amino acids) and the transmembrane domain (5 amino acids) of the E protein [29]. A similar in-frame deletion of 24 nucleotides (from position 26259 to 26284) was observed by Finkel *et al.* [28] that resulted in the loss of a subset of 8 out of the 9 amino acids in the deletion that we observed. Furthermore, it is possible that this common deletion is being selected for during passage in Vero E6 cells, which were used by both Kim *et al.* [3] and Finkel *et al.* [28]. Non-canonical transcripts S’, 3a’ and E’ also contain the same discontinuous edge from position 26256 to 26284. While transcript E’ produces a version of protein E with 9 missing amino acids, transcripts S’ and 3a’ produce complete viral proteins S and 3a, respectively. Non-canonical transcript 6’ differs from the canonical transcript 6, containing a jump from position 27886 to 27909. This jump is downstream of ORF6 and therefore does not disrupt the translation of accessory protein 6. Similarly, transcript 1ab’ has a single jump from position 26779 to 26817, which is downstream of the ORF1ab gene and therefore will yield the complete polypeptide 1ab. Transcript 7b*, on the other hand, has a single discontinuous edge from position 71 to 27762. The start codon ‘ATG’ downstream of the 3’ end occurs at position 27825, maintaining the frame of 7b, and thus leading to an N-terminal truncation [3] of 23 amino acids. Interestingly, transcript 7b and transcript 7b* appear with similar abundances in our solution. Finally, transcript N’ has one canonical discontinuous edge from TRS-L (position 65) to the transcription regulating body sequence (TRS-B) region corresponding to ORF N (position 28255) and an additional jump from position 28525 to 28577, which leads to an in-frame deletion of 17 amino acids in the N-terminal RNA-binding domain [30, 31] of ORF N. Thus, with the exception of transcripts X and X’, the non-canonical transcripts identified by Jumper either produce complete viral proteins (1ab’, S’, 3a’, 6’), contain in-frame deletions in the middle of known proteins (E’, N’) or produce N-terminally truncated proteins (7b*).

One of the major findings of the Kim *et al.* paper [3] is that the SARS-CoV-2 transcriptome is highly complex owing to numerous non-canonical discontinuous transcription events. Strikingly, our results show that these non-canonical transcription events do not significantly change the resulting proteins. Indeed, we find that 4 out of the 9 non-canonical transcripts produce a complete known viral protein and the total abundance of the predicted transcripts that produce a complete known viral protein is 0.968. Moreover, these predicted transcripts account for more than 90% of the reads in the sample according to the estimates provided by Salmon.

Typically, reads from short-read sequencing samples are not long enough to contain more than one discontinuous edge. As a result, short-read data can only provide direct evidence for transcripts with closely spaced discontinuous edges. For instance, we observe ample support (63485 short reads) for the predicted non-canonical transcript E’, which has two discontinuous edges (69, 26237) and (26256, 26284), in short-read data due to the close proximity of the two discontinuous edges (*i.e.* the discontinuous edges are only 26256 − 26237 = 19 nucleotides apart). The other non-canonical transcripts with multiple discontinuous edges, *i.e.* X’, S’, 3a’, 6’ and N’, have edges that are too far apart to be spanned by a single short read. Using the long-read sequencing data of this sample, we detect supporting long reads that span the exact set of discontinuous edges of all 9 non-canonical transcripts (Fig. 4c). Moreover, we find support for the canonical transcripts as well (Fig. C8). Thus, all transcripts identified by Jumper from the short-read data are supported by direct evidence in the long-read data.

In summary, using Jumper we reconstructed a detailed picture of the transcriptome of a short-read sequencing sample of Vero cells infected by SARS-CoV-2. While existing methods failed to recall even the reference transcriptome, Jumper identified transcripts encoding for all known viral protein products. In addition, our method predicted non-canonical transcripts and their abundances, whose presence we subsequently validated on a long-read sequencing sample of the same cells.

### 5.3 Transcript assembly in SARS-CoV-1

To show the generalizability of our method, we consider another coronavirus, SARS-CoV-1. We analyzed two published samples of human Calu-3 cells infected with SARS-CoV-1 [14], SRR1942956 and SRR1942957, with a median depth of 21,358 and 20,991, respectively. These two samples originate from the same SRA project (‘PRJNA279442’) whose metadata states that both samples were sequenced 24 hours after infection. We used `fastp` to trim the short reads (trimming parameter set to 10 nucleotides) and we aligned the resulting reads using `STAR` in two-pass mode. We ran Jumper with the 35 most abundant discontinuous edges in the segment graph. Similarly to the previous analysis, we restrict our attention to transcripts identified by Jumper that have more than 0.001 abundance as estimated by Salmon. There are 13 out of 25 such transcripts for sample SRR1942956 and 13 out of 26 such transcripts for sample SRR1942957 (Fig. 5).

**Figure 5:**
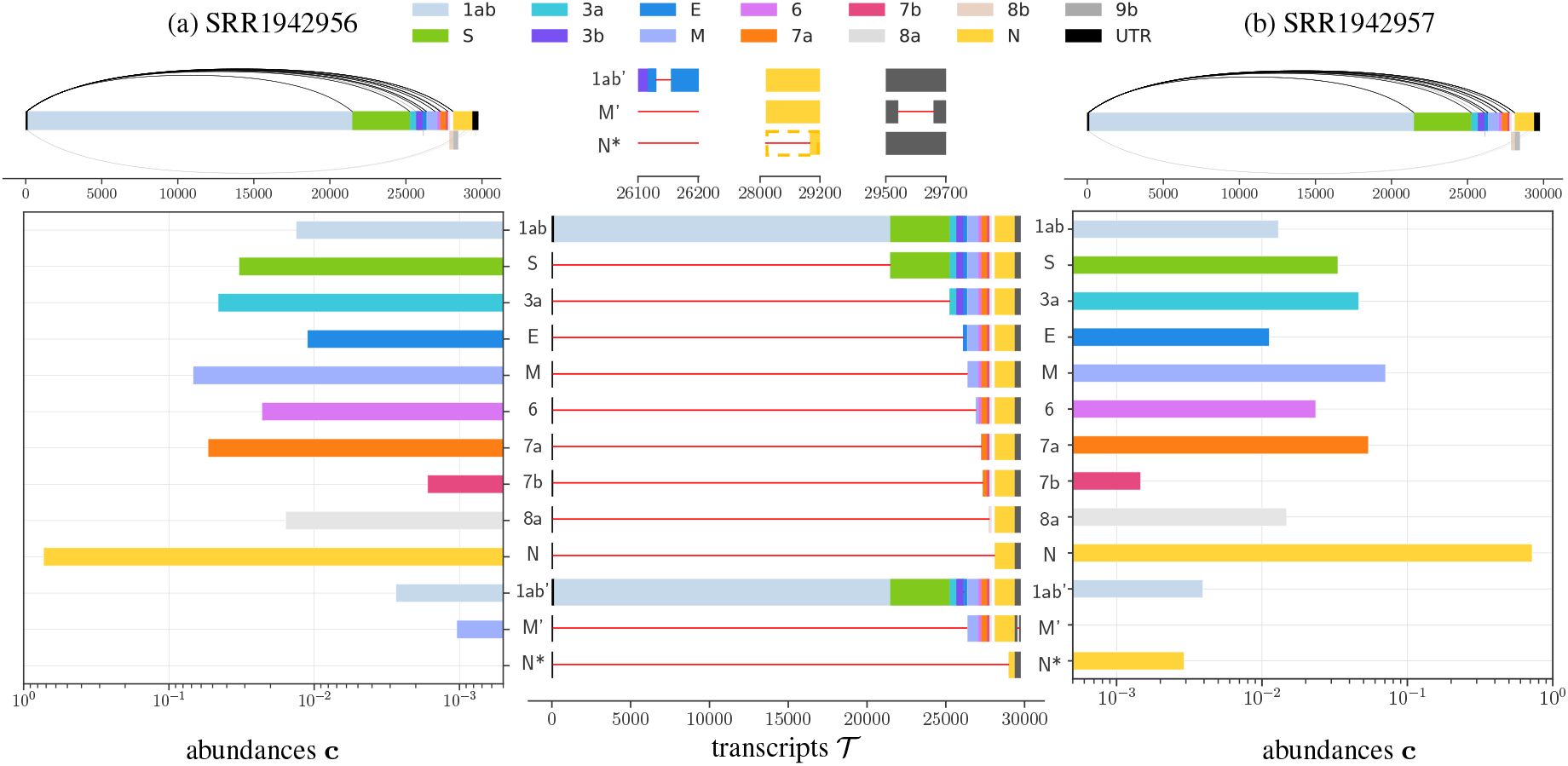
Using short-read data of SARS-CoV-1 infected Calu-3 cells [14], Jumper identifies canonical and non-canonical transcripts consistently across two samples, (a) SRR1942956 and (b) SRR1942957. For both the samples, we show the segment graph, with canonical (above) and non-canonical (below) discontinuous edges. We also show the predicted transcripts and their abundances in the two samples with a zoomed-in view of the non-canonical transcripts 1ab’, M’ and N*. UTR: untranslated region.

SARS-CoV-1 has a genome of length 29751 bp, and consists of 13 ORFs (1ab, S, 3a, 3b, E, M, 6, 7a, 7b, 8a, 8b, N and 9b), two more than SARS-CoV-2. For both samples, Jumper identifies canonical transcripts corresponding to all the ORFs of SARS-CoV-1 except ORF3b, ORF8b and ORF9b (Fig. 5). Notably, ORF8b and ORF9b share transcription regulating body sequences (TRS-B) with ORF8a and ORF N respectively [32]. More specifically, ORF9b (from position 28130 to 28426) is nested within ORF N (from position 28120 to 29388) with start codons only 10 nucleotides apart and consequently shares the same TRS-B as ORF N. ORF8b (from position 27864 to 28118) intersects with ORF8a (from 27779 to 27898) and previous studies have failed to validate a TRS-B region for ORF8b [32]. One possible way that these ORFs are translated is due to ribosome leaky scanning, which was also hypothesized to lead to ORF7b translation in SARS-CoV-2 [28]. This explains why Jumper was unable to identify transcripts that directly encode for 8b and 9b. As for ORF3b, Jumper did identify a canonical transcript corresponding to 3b in both samples, but the Salmon estimated abundances for these transcripts were below the cut-off value (0.00044 for SRR1942956 and 0.0005 for SRR1942957). Finally, we note that the relative abundances of the canonical transcripts are consistent for the two samples (Fig. 5), ranked in the same order (Fig. C9), with ORF7b being the least abundant and ORF N having the largest abundance, in line with the observations in SARS-CoV-2 (see Section 5.2).

Fig. 5 shows the three non-canonical transcripts predicted by Jumper in the two SARS-CoV-1 samples, designated as 1ab’, M’ and N*. Since these non-canonical transcripts are in very low abundance, we see some discrepancy in the prediction between the two samples. The first non-canonical transcript 1ab’ with a single short discontinuous edge from position 26131 to 26156 is detected in both samples and has a very low abundance compared to the canonical transcript 1ab (0.0133 vs 0.002 in SRR1942956, and 0.013 vs 0.0039 in SRR1942956). Since the discontinuous edge occurs downstream of the stop codon of 1ab (position 21492), the 1ab’ transcript encodes for the complete polypeptide 1ab. The second non-canonical transcript M’ has two discontinuous edges: a canonical discontinuous edge from TRS-L (position 65) to TRS-B of ORF M (position 26351) and a non-canonical discontinuous edge from 29542 to 29661 in the 3’ untranslated region (UTR). As such, this transcript encodes for the complete M protein. This transcript is detected in SRR1942956 with a very low abundance of 0.001 and is detected at an even lower abundance of 0.0008 in SRR1942957, which is below the cut-off threshold of 0.001. The third non-canonical transcript, denoted by N*, has a single discontinuous edge from position 65 to 29003. While Jumper and Salmon detect this transcript only in sample SRR1942957 with a low abundance of 0.003, we do observe 119 reads in SRR1942956 (compared to 151 reads in SRR1942957) that support this edge, suggesting that N* might be present in the latter sample at too small of an abundance to be detected. Transcript N* is interesting because the first ‘ATG’ downstream of the 3’ end of its discontinuous edge occurs at position 29071 maintaining the frame of N (which starts at position 28120). Thus transcript B* encodes for an N-terminally truncated version of protein N with 105 out the 422 amino acids that only contain part of the C-terminal dimerization domain [30]. This is similar to transcript 7b* in the SARS-CoV-2 sample (Section 5.2) which yields a N-terminal truncated version of protein 7b. Detection of non-canonical transcripts such as E’ and 7b* in SARS-CoV-2 and N’ in SARS-CoV-1 suggests that generation of N-terminally truncated proteins might be a common feature in coronaviruses. In summary, Jumper can be applied to all coronaviruses to reconstruct the transcriptome from short-read sequencing data and be used to discover novel transcripts and the viral protein products that they encode.

## 6 Discussion

In this paper, we formulated the Discontinuous Transcript Assembly (DTA) problem of reconstructing viral transcripts from short-read RNA-seq data for SARS-CoV-2 and, more generally, viruses in the order of *Nidovirales*. The discontinuous transcription process exhibited by the viral RNA-dependent RNA polymerase (RdRp) is distinct from alternative splicing observed in eukaryotes. Our proposed method, Jumper, is specifically designed to reconstruct the viral transcripts generated by discontinuous transcription and is therefore able to outperform existing transcript assembly methods such as Scallop and StringTie, as we have shown in both simulated and real data. For real-data analysis, we used publicly available short-read and long-read sequencing data of the same sample of virus infected Vero cells [3]. We performed transcript assembly using the short-read sequencing data and used the long-read data for validation. Jumper was able to identify transcripts encoding for all known viral proteins except ORF10, which has been shown to have little support of active transcription in previous studies [3, 28]. Moreover, we predicted 9 non-canonical transcripts that are well supported by long-read sequencing data. For two samples of Calu-3 cells infected by SARS-CoV-1 [14], Jumper reconstructed all the canonical transcripts with distinct TRS-B regions and additionally predicted the presence of non-canonical transcripts encoding for either complete or truncated versions of known viral proteins.

There are several avenues for future work. First, the computational complexity of the DTA problem remains open. Second, we plan to extend our current model to account for positional and sequencing biases in the data. Doing so will enable us to assemble transcriptomes from sequencing samples that used SARS-CoV-2-specific primers, which form the majority of currently available data. Third, we currently make the assumption that the read fragments are much smaller compared to the viral transcripts. We will relax this assumption in order to support long-read sequencing data that have variable read length. Fourth, we plan to study the effect of mutations (including single-nucleotide variants as well as indels) on the transcriptome. Along the same lines, there is evidence of within-host diversity in COVID-19 patients [33–38]. It will be interesting to study whether this diversity translates to distinct sets of transcripts and abundances within the same host. Fifth, there are possibly multiple optimal solutions to the DTA problem that present equally likely viral transcripts with different relative abundances in the sample. A useful direction of future work is to explore the space of optimal solutions similar to the work done in [18]. Finally, the approach presented in this paper can extended to the general transcript assembly problem, where a topological ordering of the nodes in the splice graph will serve the same function as the unique Hamiltonian path of the segment graph did in the DTA problem. We envision this will facilitate efficient use of combinatorial optimization tools such as integer linear programming to transcript assembly problems.

## Data availability

Simulated data is available at https://doi.org/10.13012/B2IDB-6667667_V1. The code is available at https://github.com/elkebir-group/Jumper.

## Acknowledgments

The authors were supported by the National Science Foundation under award number CCF-2027669.

## A Likelihood Model for Discontinuous Transcript Assembly

We use the segment graph *G* to compute the probability 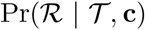 of observing the alignment 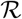 given transcripts 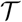 and abundances **c**. We follow the generative model described in [39], which has been extensively used for transcription quantification [19–21]. Let the set 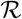 of reads be {1, … , *r_n_*} and the set 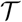 of transcripts be 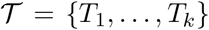 with lengths *L*_1_, … , *L_k_* and abundances **c** = [*c*_1_, … , *c_k_*]. In line with current literature, reads 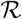 are generated independently from transcripts 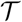 with abundances **c**. Further, we must marginalize over the set of transcripts 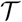 as the transcript of origin of any given read is typically unknown, due to *ℓ* ≪ *L*. Thus,

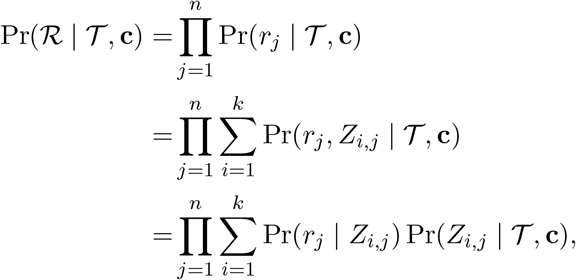

where *Z_i,j_* is the indicator random variable for the event that *T_i_* is the transcript of origin for read *r_j_*. We denote by Pr(*r_j_* | *Z_i,j_*) the probability of observing read *r_j_* given that it is generated from transcript *T_i_* and 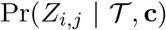 denotes the probability of generating a read from transcript *T_i_* given transcripts 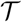 and abundances **c**.

Assuming no amplification and sequencing bias, the probability 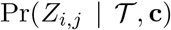 of generating a read from a transcript *T_i_* of length *L_i_* is given by

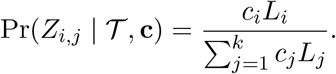

We now derive the probability Pr(*r_j_* | *Z_i,j_*) of transcript *T_i_* generating read *r_j_* of fixed length *ℓ*. We do so using the segment graph *G* = (*V, E*). Recall that a transcript *T* must correspond to an **s** to **t** path in *G*. Let *π*(*T*) ⊆ *E* denote the path corresponding to transcript *T*. Similarly, each read *r* induces a path *π*(*r*) ⊆ *E* in *G*. Read *r* can only be generated by transcript *T* if *π*(*r*) ⊆ *π*(*T*). Hence, the probability of transcript *T_i_* generating a given read *r_j_* is given by

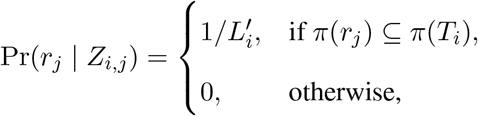

where 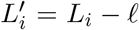 is the *effective length* of the transcript. We assume that the transcripts are much longer than the reads and as such 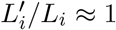. Putting it all together we get

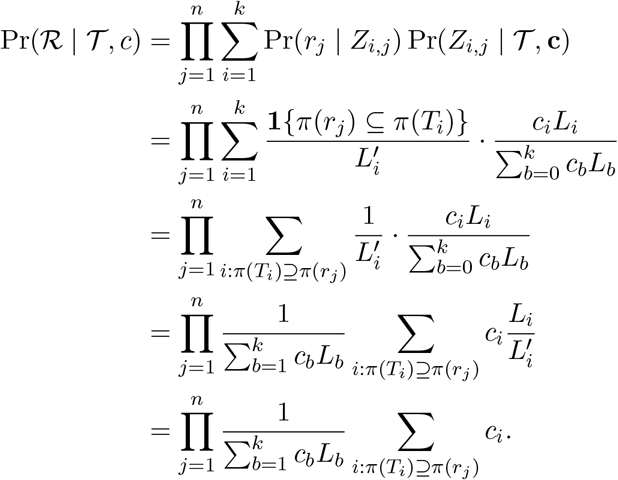

## B Supplementary Methods

## B.1 Recharacterization of solutions using discontinuous edges

We prove the following two main text propositions.

**(Main Text) Proposition 1.** There is a bijection between subsets of discontinuous edges that are pairwise non-overlapping and **s** − **t** paths in *G*.

*Proof.* Let Π be the set of **s** − **t** paths in *G*. We indicate with Σ the family of subsets of discontinuous edges that are pairwise non-overlapping. Note that 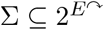.

For an **s** − **t** path *π* ∈ Π, let *f* (*π*) be the set of discontinuous edges in *π*, *i.e. f* (*π*) = *π* ∩ *E^↷^*. Since *π* is an **s** − **t** path of *G*, we have that for each edge (**v** = [*v*^−^, *v*^+^], **w** = [*w*^−^*, w*^+^]) ∈ *π* it holds that *v*^+^ ≤ *w*^−^. Therefore, *f* (*π*) is composed of pairwise non-overlapping disconnected edges.

Now, consider a subset *σ* ∈ Σ of discontinuous edges that are pairwise non-overlapping. We obtain the corresponding **s** − **t** path *f*^−1^(*σ*) by first ordering the edges of *σ* in ascending order. That is, let 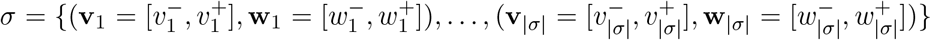 such that 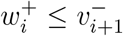 for all *i* ∈ {1, … , |*σ*|−1}. For every two consecutive discontinuous edges 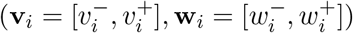 and 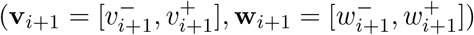, we include the corresponding subpath of continuous edges from **w**_*i*_ to **v**_*i*+1_ into *f*^−1^(*σ*). In addition, we include the subpath of continuous edges from node **s** to node **v**_1_ as well as the subpath from node **w**_|*σ*|_ to **t** into *f*^−1^(*σ*). By construction, *f*^−1^(*σ*) is an **s** − **t** path.

**(Main Text) Proposition 2.** Let *G* be a segment graph, *T* be a transcript and *r* be a read. Then, *π*(*T*) ⊇ *π*(*r*) if and only if *σ*(*T*) ⊇ *σ*^⊕^(*r*) and *σ*(*T*) ∩ *σ*^⊖^(*r*) = ∅.

*Proof.* **(⇒)** By the premise, *π*(*T*) ⊇ *π*(*r*). By definition, *σ*(*T*) = *π*(*T*) ∩ *E^↷^*. By Definition 4, *σ*^⊕^(*r*) = *π*(*r*) ∩ *E^↷^* As *π*(*T*) ⊇ *π*(*r*), we have that *σ*(*T*) = *π*(*T*) ∩ *E^↷^* ⊇ *π*(*r*) ∩ *E^↷^* = *σ*^⊕^(*r*). By definition, *σ*^⊖^(*r*) is the subset of discontinuous edges in *E^↷^* \ *σ*^⊕^(*r*) that overlaps with an edge in *π*(*r*). Since *π*(*T*) ⊇ *π*(*r*), every edge included in *σ*^⊖^(*r*) because of an overlap with an edge in *π*(*r*) must also overlap with the same edge in *π*(*T*. Since *π*(*T*) is an **s** − **t** path, and thus does not contain pairwise overlapping edges, we infer that *σ*^⊖^(*r*) ∩ *σ*(*T*) = ∅.

**(⇐)** By the premise, *σ*(*T*) ⊇ *σ*^⊕^(*r*) and *σ*(*T*) ∩ *σ*^⊖^(*r*) = ∅. As *σ*(*T*) ⊇ *σ*^⊕^(*r*), we have that *π*(*T*) ∩ *E* = *σ*(*T*) ⊇ *σ*^⊕^(*r*) = *π*(*r*) ∩ *E^↷^*. Since *σ*(*T*) ∩ *σ*^⊖^(*r*) = ∅, we have by Definition 4, that no discontinuous edge in *σ*(*T*) overlaps with any edge in *π*(*r*). Since *π*(*T*) is an **s** − **t** path containing the subset *σ*^⊕^(*r*) of discontinuous edges in *π*(*r*), it holds that *π*(*T*) ∩ *E*^→^ ⊇ *π*(*r*) ∩ *E*^→^. Finally, as *E^↷^* ∪ *E*^→^ = *E*, *π*(*r*) ⊆ *E* and *π*(*T*) ⊆ *E*, we get *π*(*T*) ⊇ *π*(*r*).

Using this proposition, we derive a simpler form of the likelihood given in Equation (2). Let 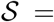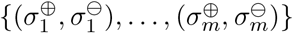 be the set of characteristic discontinuous edges generated by the reads in alignment 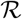. Let **d** = {*d*_1_, · · · , *d_m_*} be the number of reads that map to each pair in 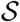. Using that distinct reads *r_j_* and *r_j′_* with the same characteristic discontinuous edges (*σ*^⊕^(*r_j_*), *σ*^⊖^(*r_j_*)) = (*σ*^⊕^(*r_j′_*), *σ*^⊖^(*r_j′_*)) have the same likelihood in terms of (2), we have

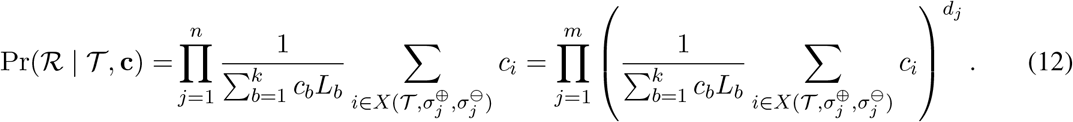

Now, taking the logarithm yields

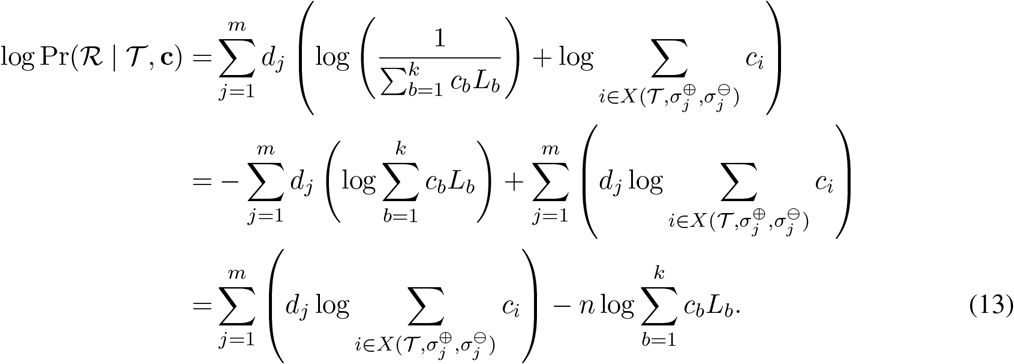

The goal is the remove the second sum in the above equation, as it is convex and we are maximizing. In order to do so, we first prove the following lemma.

### Lemma 2.

For any given scaling factor *α* > 0, we have that 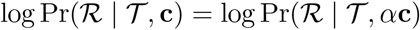.

*Proof*.

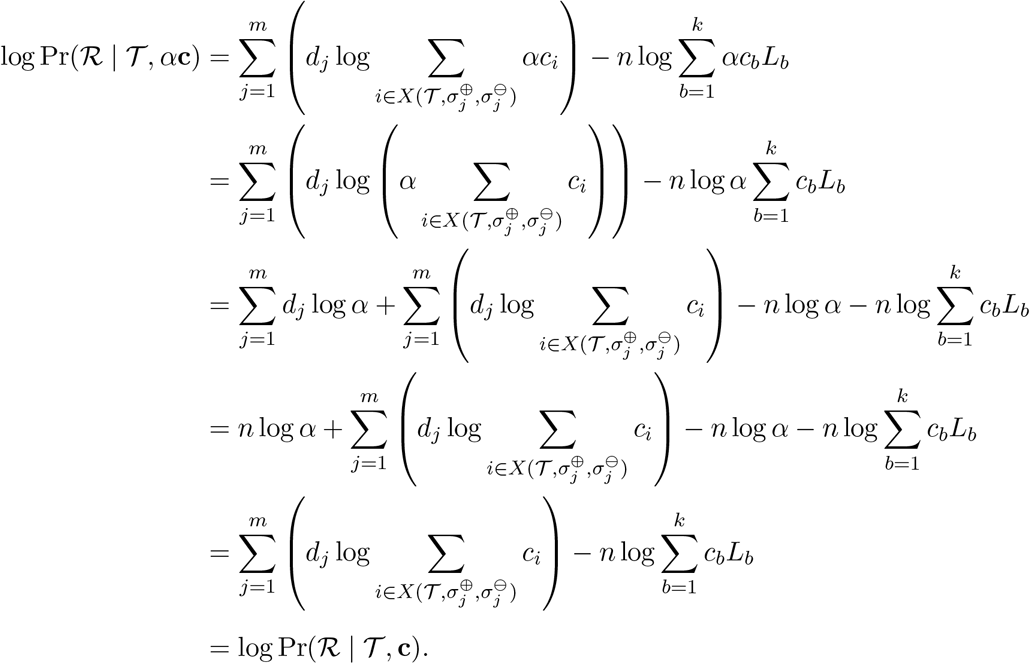

This enables us to prove the following lemma.

**(Main Text) Lemma 1.** Let *D* > 0 be a constant, 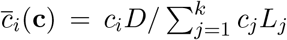 and 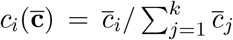 for all *i* ∈ [*k*]. Then, 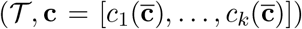 is an optimal solution for (3)-(6) if and only if 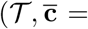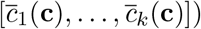 is an optimal solution for

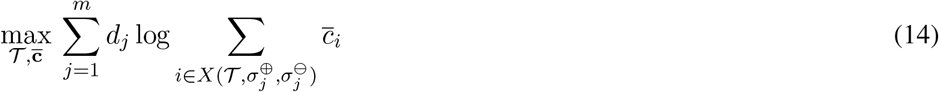

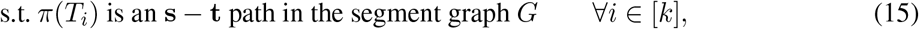

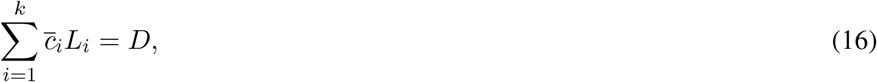

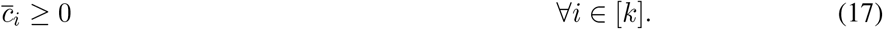

*Proof.* We will refer to the optimization problem in (3)-(6) as *P* and the optimization problem (14)-(17) as *Q*. Further, we will refer to the objective function in (3) as 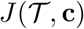 and the objective function in (14) as 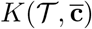. Observe that

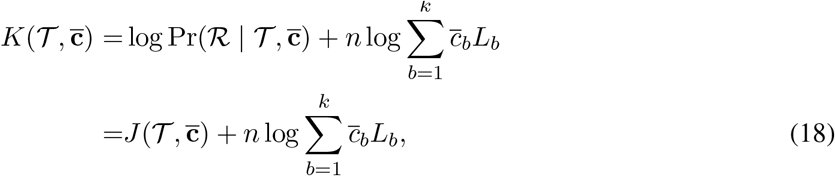

where the last equality uses (13).

**(⇒)** Let 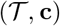 be an optimal solution to problem *P*. We begin by showing that 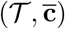 is a feasible solution to *Q* where 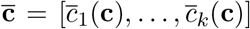. By definition of 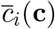, constraints (16) are satisfied. Hence, 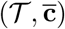 is a feasible solution to problem *Q*.

We now show that if 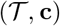 is an optimal solution to problem *P*, then 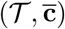 is an optimal solution to problem *Q*. Let 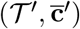 be an optimal solution to problem *Q*. Then, by optimality of 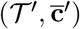, we have

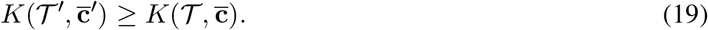

Let 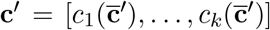. Note that **c**′ satisfies constraints (5). Thus 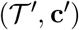 is a feasible solution to problem *P*. Since 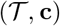 is an optimal solution of *P*, we have

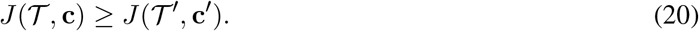

Since **c**′ and 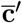 only differ by a positive scaling factor 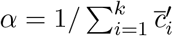, we use Lemma 2 to get 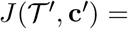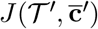. Similar result holds for **c** and 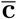, *i.e.* 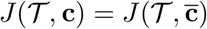. Applying this to (20), we get

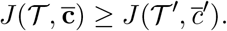

Using (16) and (18), we get

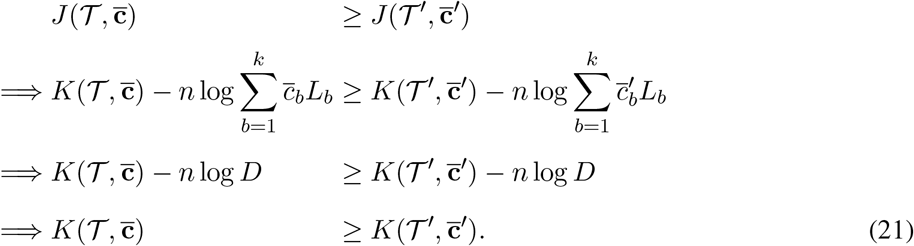

Finally, using (19) and (21), we get 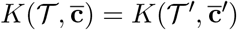. Hence, 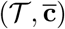 is an optimal solution of *Q*. **(⇐)** Let 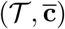 be an optimal solution to problem *Q*. We begin by showing that 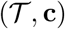 is a feasible solution to *P* where 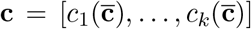. By definition of 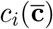, constraints (5) are satisfied. Hence, 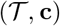 is a feasible solution to problem *P*.

Next, we need to show that 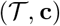 is an optimal solution to problem *P*. Let 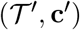 be an optimal solution to problem *P*.

Then, from the optimality condition, we get

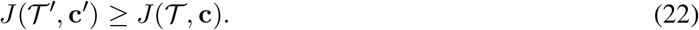

Let 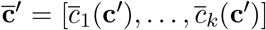. Note that 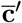 satisties constraint (16) and thus 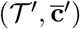 is a feasible solution to problem *Q*. Using (18) and the fact that 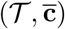 is an optimal solution of problem 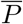 we get

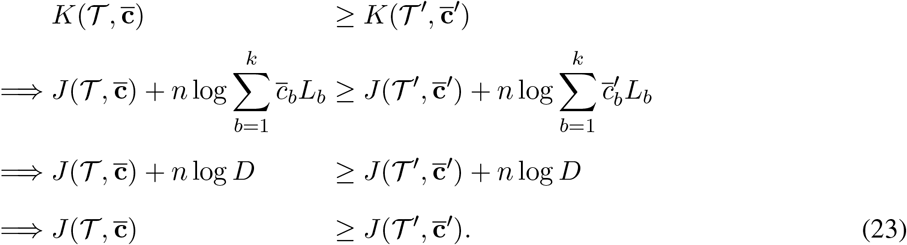

Observe that **c**′ and 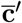 only differ by a positive scaling factor 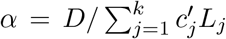. Therefore, using Lemma 2, we have 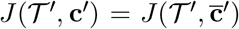. Similarly, for **c** and 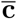, we have 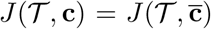. Using this together with (23), we obtain

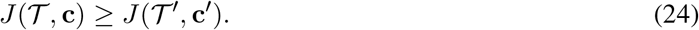

Moreover, (22) and (24) simultaneously imply 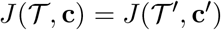. Hence, 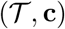 is an optimal solution to problem *P*.

## B.2 Mixed integer linear program

In the following, we introduce variables and constraints to encode the following.

i. The composition of each transcript *T_i_* as a set *σ*(*T_i_*) of non-overlapping discontinuous edges.
ii. The abundance *c_i_* and length *L_i_* of each transcript *T_i_*.
iii. The total abundance 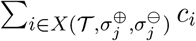 of transcripts supported by characteristic discontinuous edges 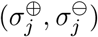.
iv. A piecewise linear approximation of the log function.

We describe (iii) and (iv) in the following and refer to Section 4 for (i) and (ii).

## Contribution of transcripts to each pair of characteristic discontinuous edges

The objective function has *m* terms, one corresponding to each pair 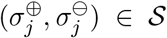 of characteristic discontinuous edges (see Eq. (7)). Specifically, each term *j* equals 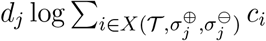 where *d_j_* is a constant, for all *j* ∈ [*m*]. We introduce non-negative continuous variables **q** = {*q*_1_, … , *q_m_*} such that

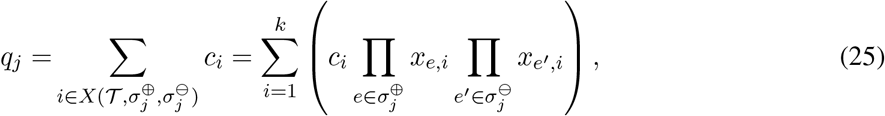

where the last equality uses the characterization of candidate transcripts of origin for a given read described in Proposition 2. We introduce continuous variables **y**_*j*_ ∈ [0, 1]^*k*^ that encode the product 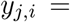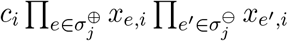. Intuitively, each variable *y_j,i_* encodes the contribution of a transcript *T_i_* for the given characteristic discontinuous edge sets 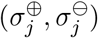. We linearize the product 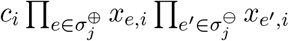 as follows.

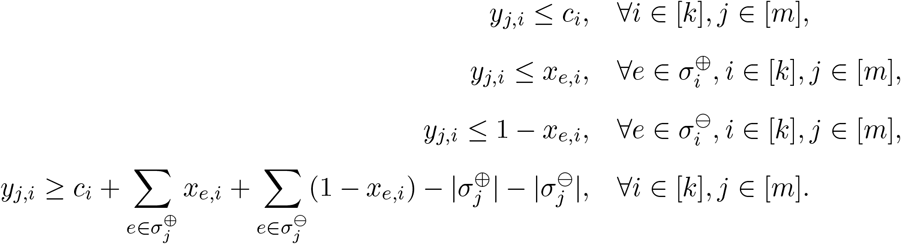

Hence, we have

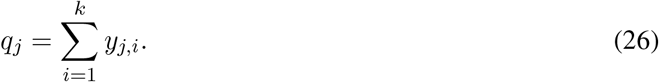

## Objective function

The objective function (Eq. (7)) can be written in terms of continuous variables **q** as

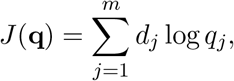

where *d_j_* is a constant and **q** is as in (26). We use the lambda method to approximate our objective method using a piecewise linear function [40]. Following the method described in [40], we partition the domain (0, 1] with *h* breakpoints *b*_1_ ≤ *b*_2_ ≤ … ≤ *b_h_*. We introduce continuous variables ***λ***_*j*_ ∈ [0, 1]^*h*^ with the constraints

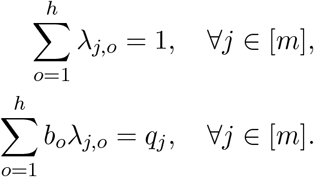

Note that *b_o_* for *o* ∈ [*h*] are constants. Since each of the *m* terms in the objective function are individually concave and we are maximizing, the adjacency condition of breakpoints does not need to be enforced. For each *j* ∈ [*m*], the log function is then approximated as

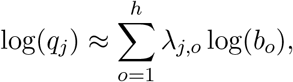

where log(*b_o_*) is a constant for each *o* ∈ [*h*]. Therefore the objective function we wish to maximize is

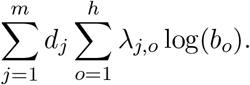

Note that since we have a log-likelihood objective function, feasibility of the solution requires that *q_j_* > 0 for *j* ∈ [*m*]. This means that for each characteristic discontinuous edge sets 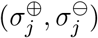, there must be at least one candidate transcript of origin *T_i_* with non-zero abundance *c_i_* > 0. This leads to the solution containing a large number of transcripts and making the problem intractable while also preventing us from finding parsimonious sets of transcripts that support most but not all of the observed reads in the sample. Finding such parsimonious solutions is often desirable since they provide a reasonable explanation of the observed reads while keeping the problem computationally tractable. In order to allow us to generate solutions that can partially explain the observed reads, we slightly modify our objective function. We introduce a new breakpoint *b*_0_ = 0 and associated continuous variables *λ*_*j*,0_ ∈ [0, 1] for *j* ∈ [*m*] so that

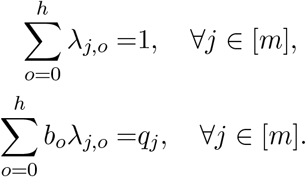

The objective function we maximize is

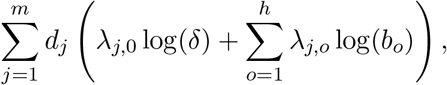

where *δ* > 0 is a small constant. Note that instead of evaluating the log function at *b*_0_, we include log(*δ*) which is well defined since *δ* > 0. In this study, we choose *δ* = *b*_1_/100 = 1/(2^*h*−1^ × 100) while *h* is left as the user’s choice with default value of 16.

Moreover, the choice of breakpoints to approximate the objective function (Eq. 7) can have a significant impact on the accuracy of the MILP solver. As a result, there has been research in efficient methods for choosing optimal breakpoint locations for convex functions, such as recursive descent algorithms [41]. In this work we take a simpler approach, by choosing breakpoints such that their spacing around a given break-point is proportional to the local gradient of the objective function. For the log function, this is equivalent to choosing breakpoints such that *b_i_* = 2^*i*−1^/2^*h*−1^. Note that *b*_0_ = 1/2^*h*−1^ while *b_h_* = 1.

## Number of variables and constraints

The total number of binary variables **x** is |*E^↷^*|*k*. Note that **q** are auxiliary (intermediate) variables that are uniquely determined by **c**, **y**, **z** and ***λ***. Therefore, the total number of required continuous variables (*i.e.* **c**, **y**, **z** and ***λ***) is *k* + *mk* + |*E^↷^*|*k* + *mh*. The number of constraints is *O*(*k*|*E*|^2^ + |*E*|*km*). We provide the full MILP for reference.

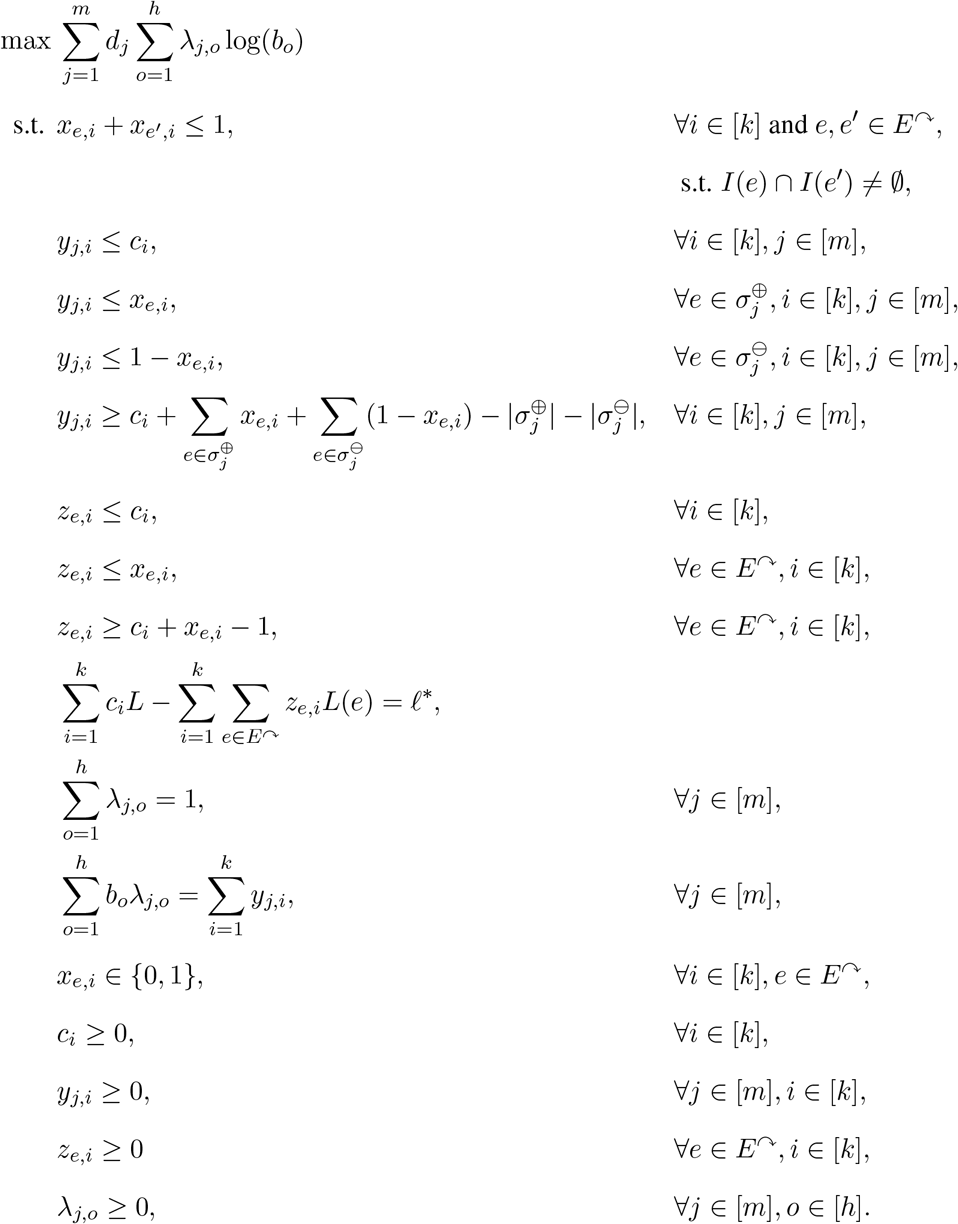

## B.3 Jumper: progressive heuristic for the DTA problem

Here we describe the subproblems that are solved at each iteration of the greedy heuristic. For a given set of transcripts 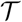 and characteristic discontinuous edge sets 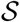, consider the optimization problem which we denote by *P*_1_,

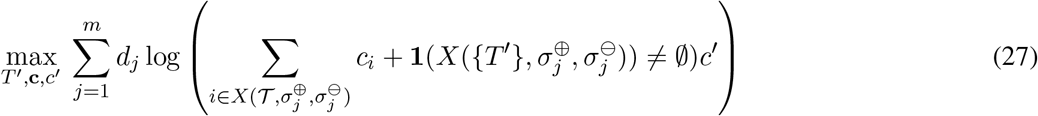

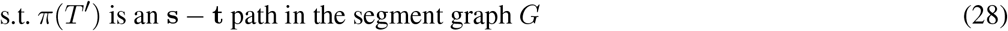

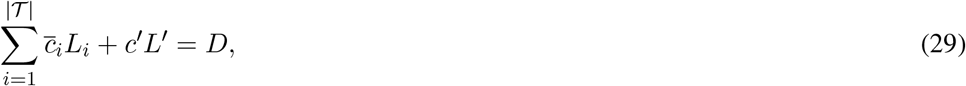

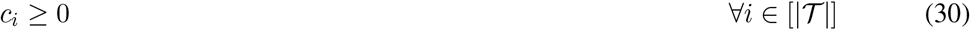

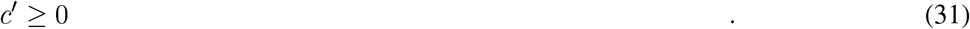

and the following optimization problem denoted by *P*_2_,

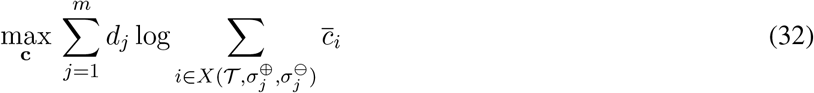

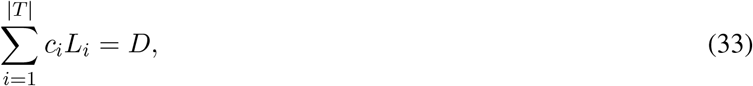

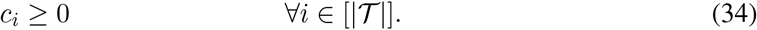

**Solution to** *P*_1_ We obtain the solution of *P*_1_ by solving the optimization problem (7)-(10) with additional constraints to fix the values of the variables that encode the presence/absence of discontinuous edges for the transcripts in 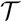. More specifically, for each transcript 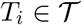, we enforce *x_e,i_* = 1 for each edge *e* ∈ *σ*(*T_i_*) and *x_e,i_* = 0 otherwise. Note that *c_i_* for 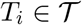 are still variables and are solved for in the optimization problem. By doing so, we only solve for the structure of the transcript *T*′ while solving for the abundance of all transcripts.

**Solution to** *P*_2_ Similar to the approach taken to solve *P*_1_, we fix the values of the variables that encode the presence/absence of discontinuous edges in the transcripts. This results in all the binary variables in the MILP with fixed values rendering the resulting optimization problem a simpler linear program.

## Heuristic Algorithm

The Algorithm 1 is re-written here in form of an itemized list.

1. Initialize 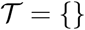, *i* = 1
2. Solve *P*_1_ with 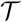 to get a new transcript *T*′ with abundance *c*′
3. Generate a new set of transcripts 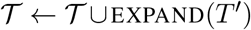 where EXPAND(*T*′) = {*T* : *σ*(*T*) ∈ 2^*σ*(*T*′)^}.
4. Solve *P*_2_ with 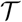 as input
5. Select *i* transcripts from 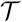. If *i* < *k* go to step (2) else return 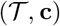

## B.4 Filtering false positive discontinuous edges

In practice, we see spurious discontinuous edges in the resulting segment graph due to sequencing and alignment errors. We filter these edges by requiring a minimum number Λ of spliced reads to support each discontinuous edge in the segment graph. The higher the value of Λ, fewer will be the number of edges and nodes in the resulting segment graph.

It is not trivial to know the right value of Λ to remove all false positive discontinuous edges. Several heuristics are used in existing methods to remove spurious splicing events. Scallop removes an edge *e* from its splice graph if the coverage of the exons of either end of the edge is more than 2*w*(*e*)^2^ + 18, where *w*(*e*) is the number of spliced reads that support the edge *e*. StringTie on the other hand, terminates its algorithm of assembling transcripts when the coverage of all the paths in the splice graph build from the un-assigned reads drops below a threshold, set by default to 2.5 reads per base-pair. By default, Jumper requires a support of 100 reads for a discontinuous edge to be included in the segment graph.

Another parameter that can be used to filter false-positive splicing events is the number of discontinuous edges allowed in the segment graph. From tests on simulated instances emulating SARS-CoV-2 samples, we found that focusing on the 35 most abundant discontinuous edges is sufficient to get a summary of the transcriptome and highly expressed canonical and non-canonical transcripts in the sample. A higher value can be used to capture more complexity of the transcriptome. By default, we set this parameter to 35.

## C Supplementary Results

## C.1 Simulation pipeline

Our simulations are based on a widely believed model of discontinuous transcription. Briefly, there are two competing models of discontinuous transcription for coronaviruses [42]. Both models agree that the RdRP jump is mediated by matching core-sequences (motifs) present in the TRSs in the viral genome. The only point of difference between the two models is whether discontinuous transcription occurs during the plus-strand synthesis or the minus-strand synthesis. The *negative-sense discontinuous transcription model* [43] proposes that the it is during the minus-strand synthesis that the RdRP performs discontinuous transcription. Transcription is initiated at the 3’ end of the plus-strand RNA and the RdRP jumps to the TRS-L region when it reaches a TRS-B region adjacent to a gene, thereby generating a minus-strand subgenomic RNA. The minus-strand subgenomic RNA is then replicated by the RdRP to produce a plus-strand RNA which can be translated to a viral protein. Currently, this model is largely believed to be true due to the considerable experimental support from genetic studies detecting minus-strand subgenomic RNAs [44–48].

We now describe the procedure to simulate transcripts and their abundances following the negative-sense model of discontinuous transcription for a given segment graph. The model is parameterized by the function *p* : *E* → [0, 1]. According to the *negative-sense discontinuous transcription* model, the transcription process is modeled as an **t** − **s** walk in the reverse graph 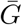 where the direction of each original edge is reversed. At each node the RdRP randomly chooses an outgoing edge to traverse in the reverse graph 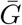 (which would be an incoming edge to the node in the original graph *G*) where the probabilities are given by the function *p*. Hence, the corresponding constraint on *p* under the negative-sense discontinuous transcription model is 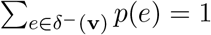. The probabilities are drawn from a Dirichlet distribution with concentration parameter *α* set to 10 for edges that are present in the path corresponding to any of the canonical transcripts and 1 otherwise. This is done to ensure that canonical transcripts are generated with high enough abundance, making the simulations similar to real data.

The next step of our simulation pipeline is to generate transcripts 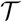 and their abundances **c** for the given segment graph. We simulate the transcription process by generating 100,000 **s** − **t** paths on the segment graph and report the number of unique paths/transcripts 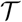 and their abundances **c**. We repeat this process to generate 5 independent sets of transcripts and abundances for the positive and the negative model each. Fig. 3b shows the number of transcripts generated from each simulation using the negative-sense discontinuous transcription model. To contrast, the total number of **s** − **t** paths in the underlying segment graph is 3440.

Once the transcripts are generated, the next step in our pipeline is to simulate the generation and sequencing of RNA-seq data. We use `polyester` [24] for this step as it allows the user to provide the number of reads generated from each transcript. For a given total number *n* of reads, the number of reads generated from transcript *T_i_* is given by 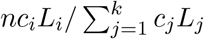 where *L_i_* is the length of the transcript *T_i_*. We use the default parameters for read length (*ℓ* = 100) and fragment length distribution (Gaussian with mean *μ_r_* = 250 and standard deviation *σ_r_* = 25) to generate 3,000,000 reads. For each set of transcript and abundances generated in the previous step of the pipeline, we simulate 5 replicates of the sequencing experiment.

The final step of the simulation pipeline is to align the generated reads to the reference genome NC 045512.2 using `STAR` [15]. The resulting BAM file serves as the input for the transcription assembly methods. To summarize, we generated 5 independent pairs 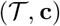 of transcripts and abundances under the negative-sense discontinuous transcription model. For each pair 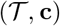 we run 5 simulated sequencing experiments using `polyester` [24]. Therefore, we generated a total of 5 × 5 = 25 simulated instances.

## C.2 Scallop arguments

We use the following arguments.

~~~
scallop -i ${input_bam} -o ${output_assembled}
~~~

## C.3 StringTie arguments

We run StringTie in de novo transcript assembly mode. That is, we do not provide a GFF file to guide assembly. We use the following arguments.

~~~
stringtie -o ${output_assembled} -A ${output_abundance} ${input_bam}
~~~

We noted that StringTie produces incomplete transcripts, *i.e.* all the assembled transcripts did not map to the 5’ and 3’ end of the reference genome. In our simulations, StringTie was not penalized for this as our evaluation metrics considered only discontinuous edges.

## C.4 Human gene simulations

We evaluate the performance of Jumper, Scallop and StringTie on simulated samples of a single human gene as well. We use a selected region (from position 5001 to 30255) of the FAS gene as the reference genome^1^ with is located on the long arm of chromosome 10 in humans and encodes the Fas cell surface receptor which leads to programmed cell death if it binds its ligand (Fas ligand).

We include the following 3 distinct isoforms of this gene (P25445-1, P25445-6 and P25445-7) with equal proportions in the ground truth.

**Figure C1:**
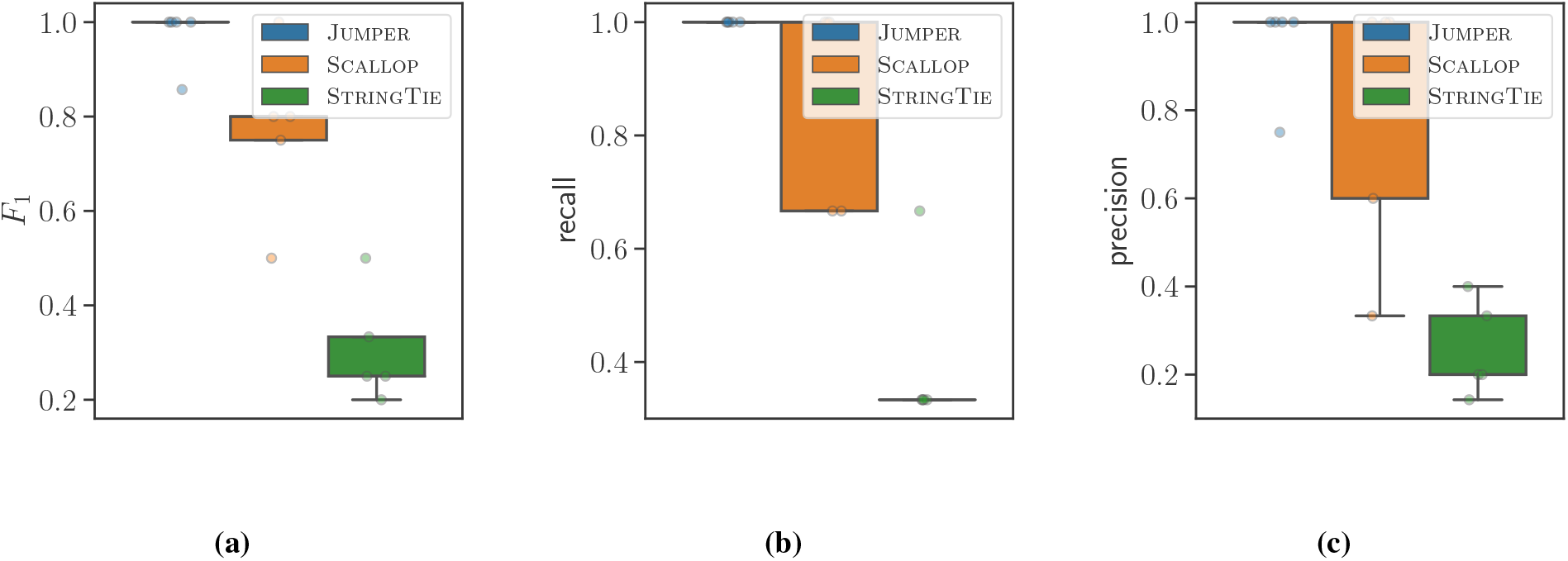
Jumper outperforms Scallop and StringTie for all simulation instances of the FAS gene (on human chromosome 10) in terms of *F*_1_ score, recall and precision while maintaining a modest running time. (a) *F*_1_ score (b) recall and (c) precision of the three methods for the simulated instances. The ground truth contained three isoforms of the FAS gene with uniform relative abundances.

1. P25445-1 with length of 335aa https://www.ncbi.nlm.nih.gov/CCDS/CcdsBrowse.cgi?REQUEST=CCDS&DATA=CCDS7393.1
2. P25445-6 with length of 314aa https://www.ncbi.nlm.nih.gov/CCDS/CcdsBrowse.cgi?REQUEST=CCDS&DATA=CCDS7394.1
3. P25445-7 with length of 220aa https://www.ncbi.nlm.nih.gov/CCDS/CcdsBrowse.cgi?REQUEST=CCDS&DATA=CCDS7395.1

We add a poly-A tail of length 85 at the end of the reference genome as well as each of the isoforms to emulate the transcription process.

We use polyester [24] to simulate the sequencing of 30,000,000 paired-end reads of the sample with a Gaussian fragment length distribution with mean 250 and standard deviation of 25. We simulate 5 replicates of the sequencing experiment. The simulated reads are aligned to the selected region of the FAS gene using `STAR` [15]. The resulting BAM file serves as the input for the transcription assembly methods. Figure C1 shows that Jumper outperforms both Scallop and StringTie in terms of both recall and precision.

## C.5 Supplementary results figures

We have the following supplementary figures.

- Fig. C2 shows that Jumper outperforms Scallop and StringTie for all simulation instances in terms of *F*_1_ score, recall and precision while maintaining a modest running time.
- Fig. C3 shows that Jumper outperforms Scallop and StringTie for varying values of thresholding parameter Λ.
- Fig. C4 shows that Jumper produces better recall and precision when compared to Scallop and StringTie for every simulation instance 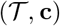.
- Fig. C5 shows that the core sequence observed in the reference genome potentially explaining a non-canonical discontinuous transcription event, and the core sequence corresponding to transcript X is conserved across *Sarbecovirus* species.
- Fig. C6 shows an example of a supporting read for a transcript with two discontinuous edges.
- Fig. C7 shows that transcript X is supported in both long-read and short-read samples deposited in SRA.
- Fig. C8 shows the number of *supporting reads* with the 5’ end mapping to the leader sequence in the short and long read sequencing data.
- Fig. C9 shows the abundances of the predicted transcripts by Jumper in two SARS-CoV-1 infected samples.
- Table C1 shows summary of the results from the simulations.

**Figure C2:**
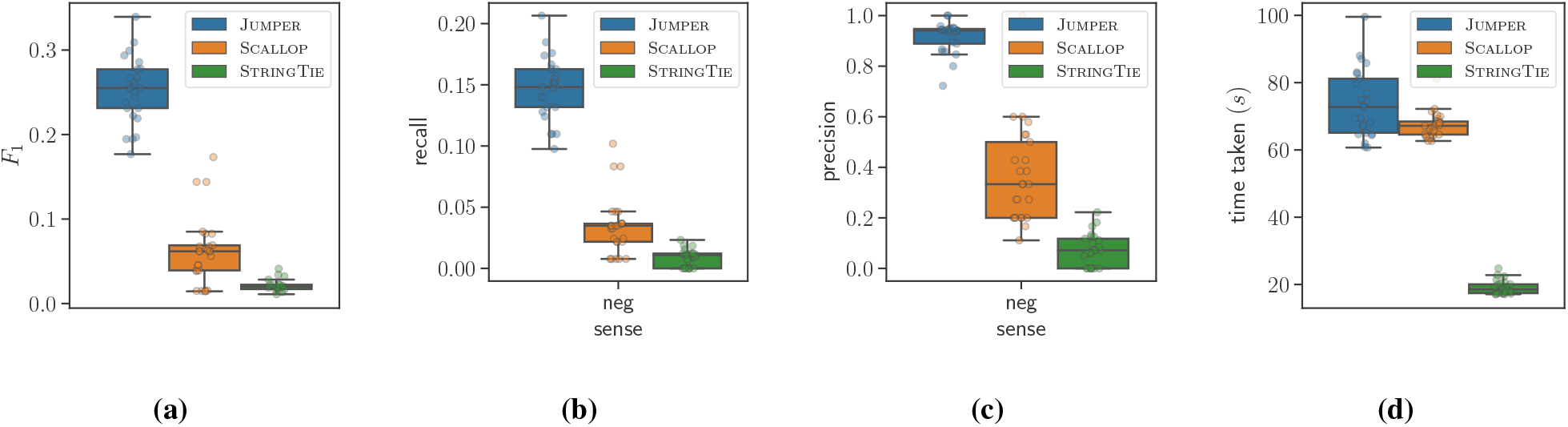
Jumper outperforms Scallop and StringTie for all simulation instances in terms of *F*_1_ score, recall and precision while maintaining a modest running time. (a) *F*_1_ score (b) recall, (c) precision and (d) time taken by the three methods for the simulated instances.

**Figure C3:**
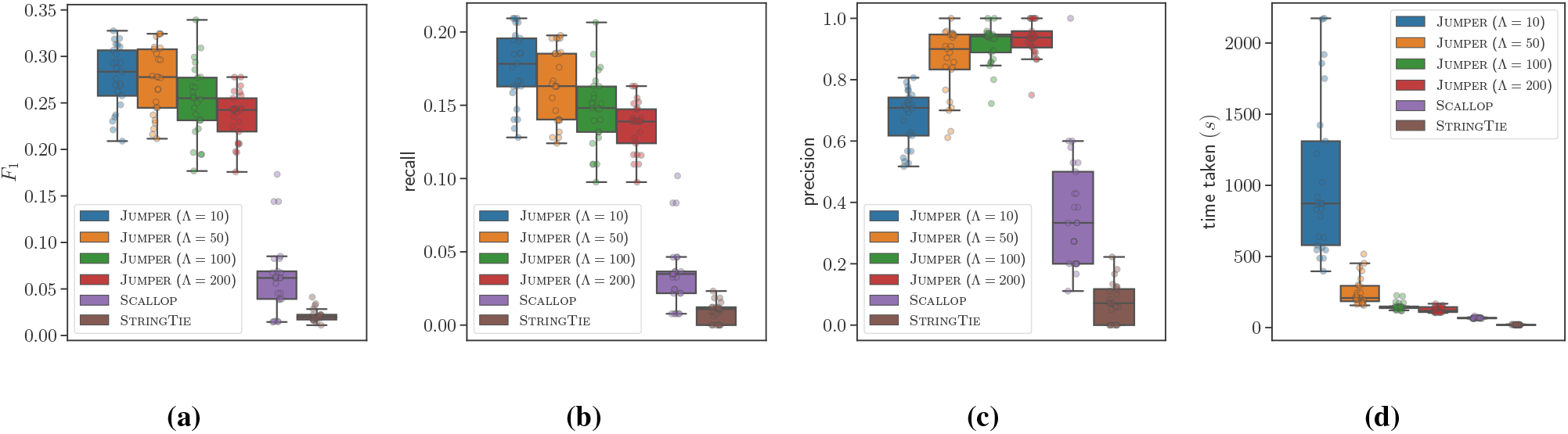
Jumper outperforms Scallop and StringTie for varying values of thresholding parameter Λ. (a) *F*_1_ score (b) recall, (c) precision and (d) time taken by the Jumper for different values of Λ compared to Scallop and StringTie on the simulated instances. As expected, the recall value drops for increasing Λ while the precision increases. We set the default value of Λ to 100 which incurs runtime comparable to Scallop while producing higher recall and precision solutions.

**Figure C4:**
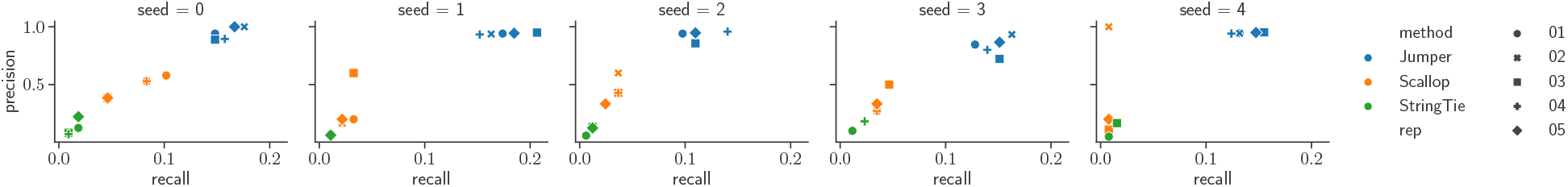
While all three methods return consistent results when generating technical sequencing replicates, Jumper produces better recall and precision when compared to Scallop and StringTie for every simulation instance 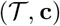. Varying simulation instances 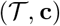 correspond to distinct panels. Each panel shows the recall and precision of the three methods for 5 sequencing experiments of the same simulated instance 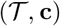.

**Figure C5:**
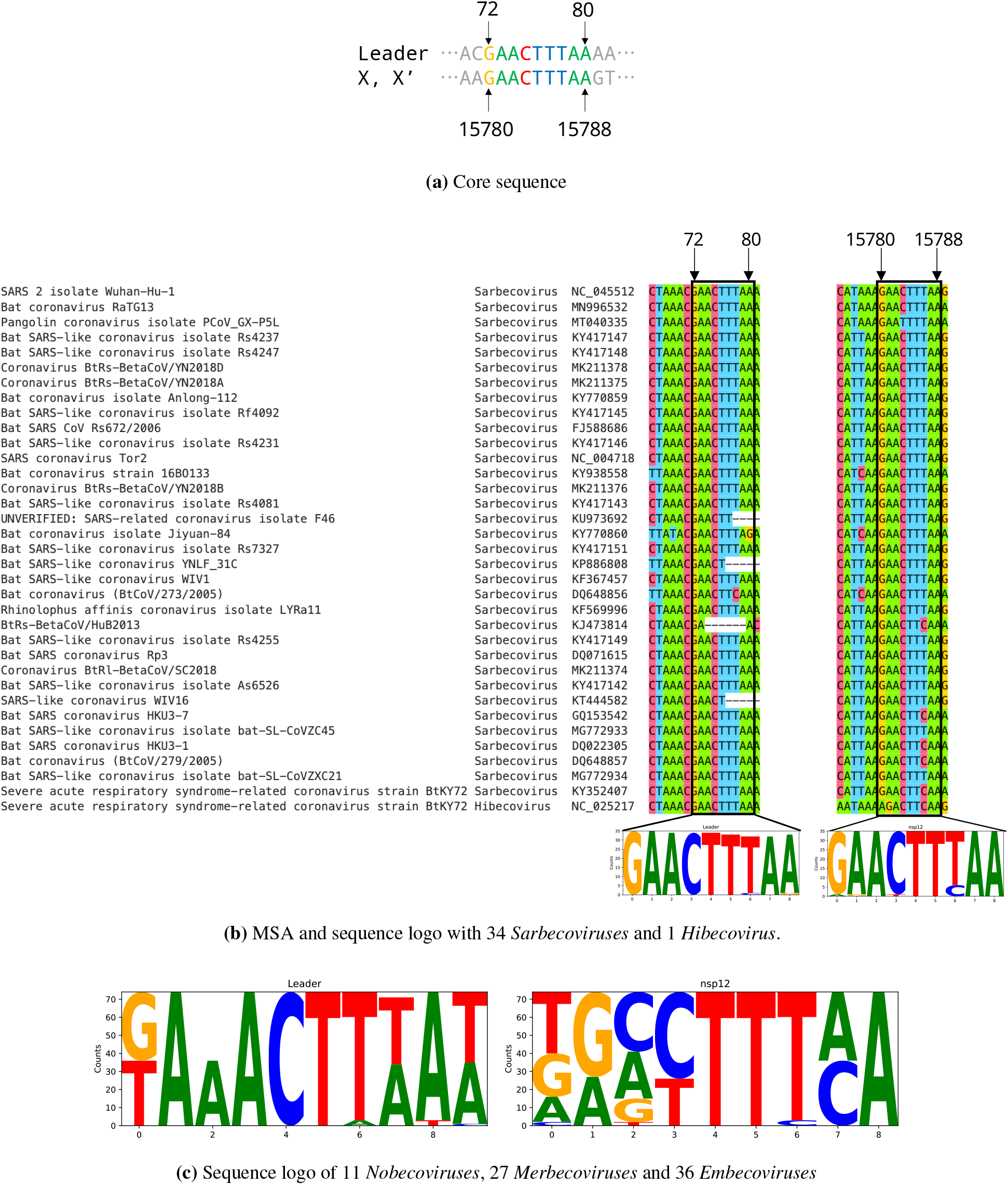
The core sequence of transcript X is conserved within the *Sarbecovirus* subgenus but not in other subgenera of the *Betacoronavirus* genus. (a) Core sequence for the transcript X and X’. (b) Sequence logo for the positions 15780 to 15788 in SARS-CoV-2 genome built from the multiple sequence alignment of the leader sequence and ORF1ab of 34 *Sarbecovirus* and a *Hibecovirus*. (c) Sequence logo for positions 15780 to 15788 in SARS-CoV-2 genome built from multiple sequence alignment with the remaining subgenera of *Betacoronaviruses*.

**Figure C6:**
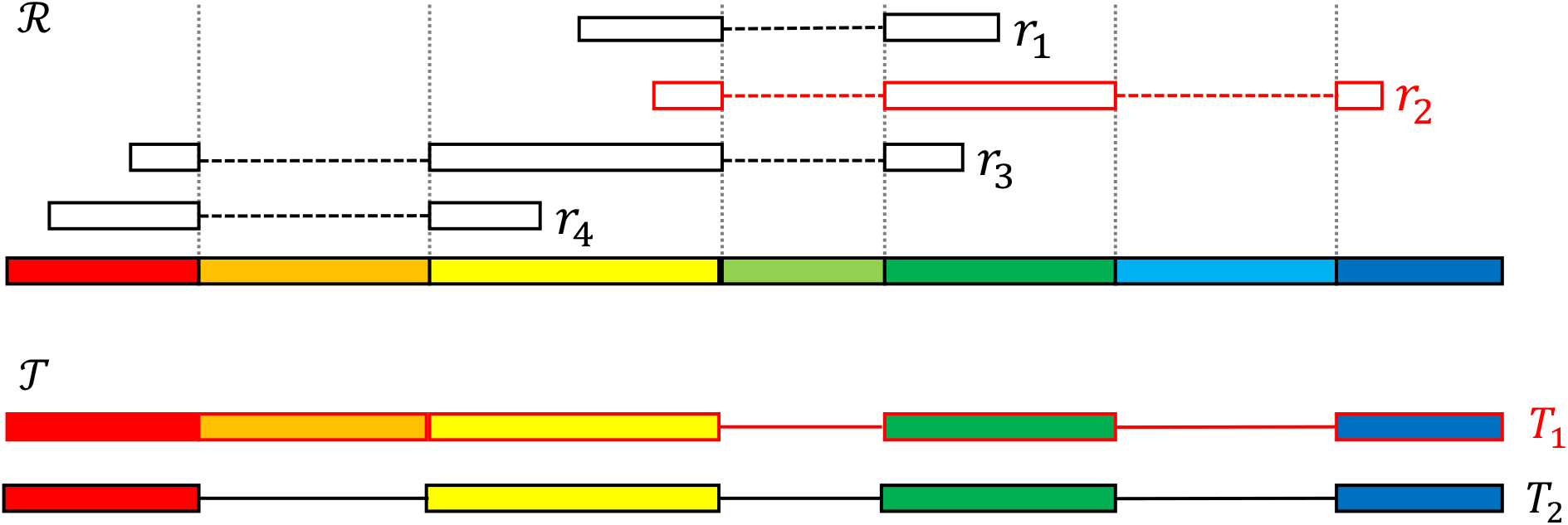
A schematic showing an example of a supporting read for a transcript *T*_1_ with *σ*^⊕^(*T*_1_) = 2. Transcript *T*_1_ is supported by *r*_2_ because *π*(*r*_2_) = *π*(*T*_1_) and |*σ*^⊕^(*r*_1_)| = |*σ*^⊕^(*T*_1_)| = 2. Reads *r*_1_*, r*_3_ and *r*_4_ do not support *T*_1_ since |*σ*^⊕^(*r*_1_)| < |*σ*^⊕^(*T*_1_)| and *π*(*r*_3_)*, π*(*r*_4_) ⊈ *π*(*T*_1_). No reads support *T*_2_ since |*σ*^⊕^(*r_j_*)| < |*σ*^⊕^(*T*_2_)| for all reads *r_j_*.

**Figure C7:**
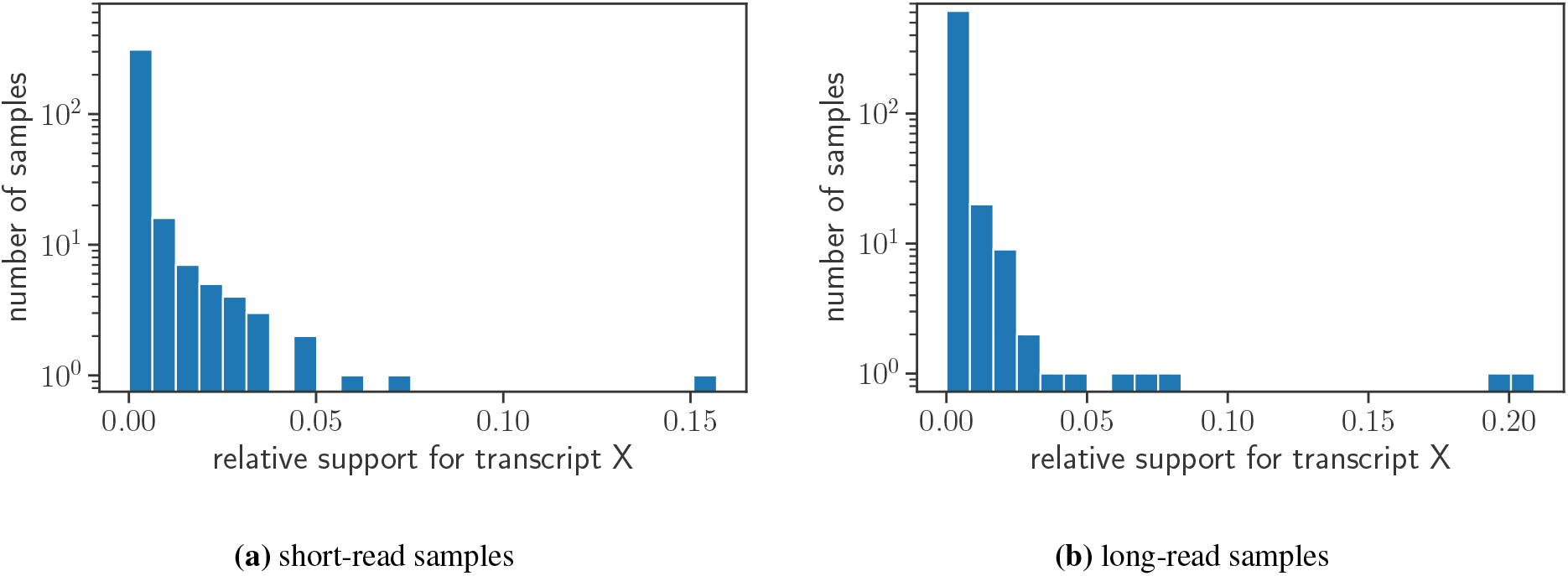
Transcript X has supporting reads in multiple independent SRA samples of SARS-CoV-2. Distribution of number of (a) short-read and (b) long-read SRA samples with varying proportion of leader-sequence spanning reads that support transcript X. All the short-read samples were aligned using `STAR` [15] while the long-read samples were aligned using `minimap2` [27]. In this plot we only consider samples with more than 100 reads that map to the leader-sequence (position 55 to 85 in the SARS-CoV-2 reference genome).

**Figure C8:**
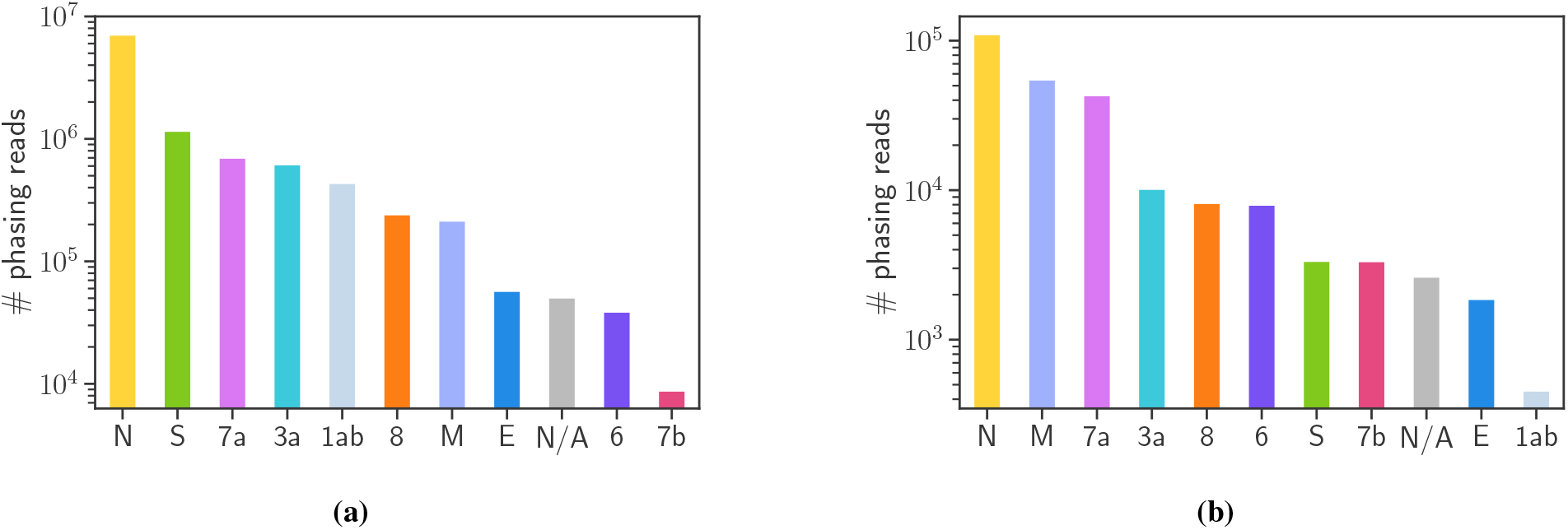
Supporting phasing reads with 5’ end mapping to the leader sequence in short and long-read sequencing samples of SARS-CoV-2 infected Vero cells [3]. Supporting phasing reads have at most one discontinuous edge with the 5’ end occurring in the leader sequence (*i.e.* between positions 55 and 85) and the first occurrence of ‘AUG’ downstream of the 3’ end position coinciding with the start codon of a known ORF. Supporting phasing reads corresponding to ‘1ab’ start in the leader sequence but do not contain a discontinuous edge. Supporting phasing reads corresponding to ‘N/A’ start in the leader sequence but have a 3’ end such that the first occurrence of ‘AUG’ down-stream of the 3’ end position does *not* coincide with the start codon of any known ORFs. (a) Supporting phasing reads in the short-read sequencing sample. (b) Supporting phasing reads in the long-read sequencing sample.

**Figure C9:**
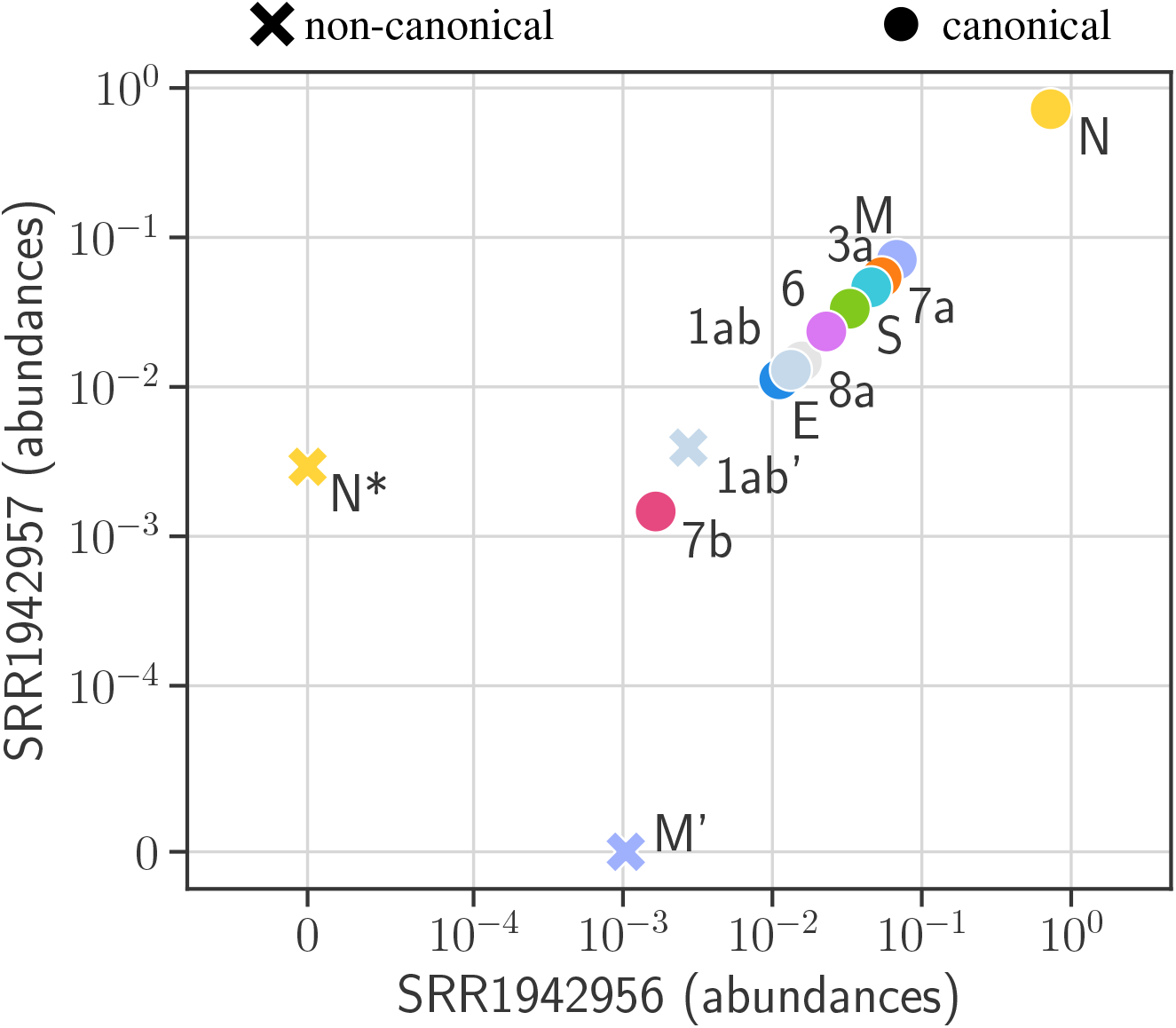
Abundances of the canonical and non-canonical transcripts predicted by Jumper are consistent in the two SARS-CoV-1 infected samples (SRR194256 and SRR194257). Jumper predicts 10 canonical and 3 non-canonical transcripts across the two samples.

**Table C1:** Simulation results for the three methods Jumper, Scallop and StringTie. Each distinct value in the column ‘seed’ is a unique instance of 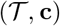 and each distinct value in the column ‘rep’ is a unique sequencing experiment for the given 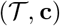. (rep: replicate, can: canonical, non-can: non-canonical, TP: true positives, FP: false positives)

| Simulation |  |  |  | JUMPER |  |  | SCALLOP |  |  | STRINGTIE |  |  |
| --- | --- | --- | --- | --- | --- | --- | --- | --- | --- | --- | --- | --- |
| seed | rep | can | non-can | TP |  | FP | TP |  | FP | TP |  | FP |
|  |  |  |  | can | non-can |  | can | non-can |  | can | non-can |  |
| 0 | 1 | 14 | 94 | 7 | 9 | 1 | 7 | 4 | 8 | 2 | 0 | 14 |
| 0 | 2 | 14 | 94 | 8 | 11 | 0 | 7 | 2 | 8 | 1 | 0 | 13 |
| 0 | 3 | 14 | 94 | 7 | 11 | 2 | 4 | 1 | 8 | 1 | 0 | 11 |
| 0 | 4 | 14 | 94 | 6 | 9 | 2 | 7 | 2 | 8 | 1 | 0 | 13 |
| 0 | 5 | 14 | 94 | 7 | 11 | 0 | 4 | 1 | 8 | 1 | 0 | 7 |
| 1 | 1 | 14 | 78 | 3 | 13 | 1 | 3 | 0 | 12 | 2 | 0 | 13 |
| 1 | 2 | 14 | 78 | 4 | 11 | 1 | 2 | 0 | 10 | 1 | 0 | 13 |
| 1 | 3 | 14 | 78 | 3 | 16 | 1 | 3 | 0 | 2 | 1 | 0 | 12 |
| 1 | 4 | 14 | 78 | 3 | 11 | 1 | 2 | 0 | 8 | 0 | 0 | 16 |
| 1 | 5 | 14 | 78 | 4 | 13 | 1 | 2 | 0 | 8 | 1 | 0 | 15 |
| 2 | 1 | 14 | 150 | 5 | 11 | 1 | 3 | 1 | 8 | 1 | 0 | 16 |
| 2 | 2 | 14 | 150 | 4 | 14 | 3 | 5 | 1 | 4 | 2 | 0 | 15 |
| 2 | 3 | 14 | 150 | 5 | 13 | 3 | 5 | 1 | 8 | 2 | 0 | 13 |
| 2 | 4 | 14 | 150 | 7 | 16 | 1 | 5 | 1 | 8 | 2 | 0 | 16 |
| 2 | 5 | 14 | 150 | 4 | 14 | 1 | 3 | 1 | 8 | 2 | 0 | 14 |
| 3 | 1 | 14 | 72 | 4 | 7 | 2 | 3 | 0 | 8 | 1 | 0 | 9 |
| 3 | 2 | 14 | 72 | 6 | 8 | 2 | 3 | 0 | 4 | 0 | 0 | 8 |
| 3 | 3 | 14 | 72 | 7 | 6 | 4 | 4 | 0 | 8 | 0 | 0 | 20 |
| 3 | 4 | 14 | 72 | 4 | 8 | 3 | 3 | 0 | 8 | 2 | 0 | 9 |
| 3 | 5 | 14 | 72 | 4 | 9 | 2 | 3 | 0 | 6 | 0 | 0 | 4 |
| 4 | 1 | 14 | 115 | 4 | 13 | 1 | 1 | 0 | 4 | 1 | 0 | 19 |
| 4 | 2 | 14 | 115 | 5 | 12 | 1 | 1 | 0 | 0 | 0 | 0 | 12 |
| 4 | 3 | 14 | 115 | 6 | 14 | 1 | 1 | 0 | 8 | 2 | 0 | 10 |
| 4 | 4 | 14 | 115 | 6 | 10 | 1 | 1 | 0 | 4 | 0 | 0 | 16 |
| 4 | 5 | 14 | 115 | 6 | 13 | 1 | 1 | 0 | 4 | 0 | 0 | 12 |

## Footnotes

1 NCBI reference sequence NG 009089.2: https://www.ncbi.nlm.nih.gov/nuccore/NG_009089.2?from=5001&to=30255&report=fasta

